# The BraALA3 Homologs Mediate Propiconazole-Modulated Plant Growth in flowering Chinese cabbage

**DOI:** 10.64898/2026.04.16.718850

**Authors:** Qing Gao, Yunxue Song, Yongsheng Yang, Siwei Wang, Xiancong Ruan, Zhicheng Liu, Dekang Guo, Yiping Chen, Xinping Wang, Ronghua Chen, Hanhong Xu, Fei Lin

**Author notes:** Corresponding Authors: **Ronghua Chen** − *Institute of Nanfan & Seed Industry, Guangdong Academy of Sciences, Guangzhou 510316, China;*; **Hanhong Xu** − *State Key Laboratory of Green Pesticide/State Key Laboratory for Conservation and Utilization of Subtropical Agro-Bioresources/Key Laboratory of Natural Pesticide and Chemical Biology, Ministry of Education, College of Plant Protection, South China Agricultural University, Guangzhou 510642, China;*; **Fei Lin** − *State Key Laboratory of Green Pesticide/State Key Laboratory for Conservation and Utilization of Subtropical Agro-Bioresources/Key Laboratory of Natural Pesticide and Chemical Biology, Ministry of Education, College of Plant Protection, South China Agricultural University, Guangzhou 510642, China;.

## Abstract

In agriculture, propiconazole (PCZ) controls excessive growth in flowering Chinese cabbage but poses dietary safety risks due to residue accumulation. Therefore, identifying novel PCZ targets and breeding PCZ-free cultivars is critical for the safe production of flowering Chinese cabbage. Here, we identified three P4-ATPase flippase homologs aminophospholipid ATPase 3 (BraALA3a/b/c) in flowering Chinese cabbage that function as sensitive targets for PCZ. These proteins exhibit high binding affinity for PCZ, which directly inhibits their ATPase activity. Overexpression of the *BraALA3* homologs enhanced plant growth and increased sensitivity to PCZ, whereas knockdown led to dwarfism and reduced sensitivity. Based on these findings, we identified editable active sites via protoplast-based screening. Genetic transformation of one such site yielded BraALA3a/*braala3a^K200T^* mutant lines, which displayed a dwarf and compact architecture. These findings provide a precise molecular target for developing PCZ-free germplasm in flowering Chinese cabbage through gene editing.

## INTRODUCTION

Flowering Chinese cabbage (*Brassica rapa* var. *parachinensis*), also known as Caixin, a leafy cruciferous vegetable of the Brassicaceae family, is characterized by emerald-green foliage, succulent stems, and a subtly sweet flavor.^1,2^ Originating in southern China, it has become a significant crop for both dietary nutrition and agricultural exports.^3,4^ Unlike heading Chinese cabbage, flowering Chinese cabbage undergoes concurrent vegetative and reproductive growth. This unique developmental pattern, coupled with abundant water and light-thermal resources in typical cultivation environments, often leads to excessive vegetative growth and premature bolting, ultimately reducing yield and quality.^5,6^ In agricultural practice, the application of propiconazole (PCZ) has been adopted to modulate plant architecture in flowering Chinese cabbage, inhibiting excessive vegetative growth, thereby aligning with consumer preference for compact and sturdy phenotypes.^7,8^ However, the brief growth period of flowering Chinese cabbage imposes a significant constraint on the safe preharvest interval for PCZ, resulting in pesticide accumulation and residues that threatening dietary safety.^9,10^ Therefore, elucidating the molecular mechanism of PCZ-mediated plant architecture regulation, identifying potential novel pesticide targets, and developing PCZ-free cultivars are of paramount importance for ensuring the safe production of flowering Chinese cabbage.

Previous studies have indicated that PCZ as a triazole fungicide primarily functions by inhibiting fungal CYP51 (14α-demethylase) activity.^11,12^ In plants, CYP51 catalyzes the initial cyclization step in sterol biosynthesis and participates in the formation of brassinosteroids.^13^ Rescue experiments with intermediates of brassinosteroid biosynthesis, combined with *in vitro* protein inhibition assays, revealed that propiconazole specifically targets the C22 and C23 side-chain hydroxylation reactions during the conversion of campesterol to teasterone.^14^ While the effect of PCZ on plant sterol and brassinosteroid biosynthesis is known,^14^ how it interacts with membrane structures upon entering cells remains unclear. It has been reported that PCZ inhibits pollen germination and tube elongation in *Tradescantia virginiana*, while disrupting cytoplasmic streaming and cytoskeletal organization. This implies that its effects may be mediated through perturbations in membrane lipid composition or integrity, which then indirectly impair cytoskeleton-membrane interactions rather than through direct targeting of microfilaments or microtubules.^15^ This suggests that transmembrane structural components may represent critical targets for PCZ response. As core regulators of membrane homeostasis, P4-ATPases are widely conserved across eukaryotes. Phospholipid flippases, classified as P4-ATPases, are integral membrane proteins that catalyze ATP-dependent translocation of phospholipids across eukaryotic membranes.^16^ Research on P4-ATPases has elucidated the profound connection between asymmetric phospholipid distribution and essential cellular functions.^17^ The research trajectory began with the 1984 discovery of phosphatidylserine (PS) transbilayer movement in human erythrocytes,^18^ leading to recognition of an ATP-dependent enzymatic system driving this process. A pivotal breakthrough came in 1996 with the identification of yeast P4-ATPase Drs2p, whose deletion impaired PS translocation from the exoplasmic to cytoplasmic leaflet.^19^ Subsequent studies revealed that Drs2p forms a functional complex with its *β*-subunit Cdc50p at the TGN, where it orchestrates clathrin-coated vesicle biogenesis through maintenance of membrane phospholipid asymmetry,^20,21^ demonstrating that P4-ATPases serve not merely as lipid “transporters” but as fundamental “architects” of membrane trafficking systems. The year 2019 marked a pivotal advance in the structural biology of P4-ATPases, with the reporting of the cryo-EM structure of the yeast Drs2p-Cdc50p complex. The structure revealed Drs2p’s ten transmembrane helices and three cytosolic domains, showing that the complex is autoinhibited by the C-terminal tail of Drs2p and activated by the lipid phosphatidylinositol 4-phosphate (PI4P) .^22^ Concurrently, cryo-EM structures of the human ATP8A1-CDC50 complex in multiple states were solved, elucidating how phosphorylation-driven conformational changes in the M1-M2 domains accomplish the transmembrane flipping of phosphatidylserine.^23^ The plant protein ALA3 (aminophospholipid ATPase 3), a homolog of yeast DRS2, localizes to the plasma membrane (PM), trans-Golgi network (TGN), and Golgi apparatus.^24^ Phenotypic analysis of the *Arabidopsis thaliana* (*A. thaliana*) *atala3* mutant revealed compromised root and shoot growth, accompanied by shorter, rounder rosette leaves.^25^ Through its ATPase-mediated phospholipid flipping activity, ALA3 is involved in coordinating the regulation of PIN protein polar trafficking. Loss-of-function mutations (*ala3/repp1*2) cause basipetal-to-acropetal polarity inversion of PIN1-HA in root epidermal cells, disrupting auxin gradient formation and resulting in gravitropic defects and aberrant lateral root development.^24^

Addressing the pressing issue of PCZ accumulation and residue in flowering Chinese cabbage production, this study elucidates the molecular mechanisms underlying PCZ-regulated plant architecture through its targeting of the membrane proteins BraALA3a/b/c. We identified three P4-ATPase flippase homologs (BraALA3a/b/c) as sensitivity factors for PCZ. *In vitro* assays demonstrated that PCZ exhibited binding affinity for BraALA3 proteins and inhibited their ATPase activity. *In vivo* assays revealed that knockdown of *BraALA3a/b/c* resulted in dwarfism and reduced sensitivity to PCZ, whereas overexpression promoted plant growth and enhanced sensitivity to PCZ. Furthermore, using a protoplast-based editing system, three editable active sites were identified in the *BraALA3a* gene. Genetic transformation yielded BraALA3a/*braala3a^K200T^* mutant lines exhibiting a dwarf and compact architecture. Collectively, these findings provide a gene-editing-based sustainable alternative for controlling excessive vegetative growth in flowering Chinese cabbage.

## MATERIALS AND METHODS

### Chemicals and Materials

PCZ was purchased from Shanghai Bepharm Science & Technology Co., Ltd. Other reagents were purchased from Sigma-Aldrich Chemical Corporation, Ltd. (St. Louis, MO, USA). Primers were synthesized by Tsingke Biotechnology Co., Ltd. (Beijing, China) (Table S2). The *Saccharomyces cerevisiae* BJ5465 strain was obtained from American Type Culture Collection (ATCC).

### Plant Materials and Growth Conditions

All flowering Chinese cabbage materials were of the ’Youqing Sijiu’ ecotype background.^26^ Genetic modifications in this background included *BraALA3a/b/c* knockdown (*BraALA3a/b/c^KD^*), gene-edited (BraALA3a/*braala3a^K200T^*), and overexpression (*BraALA3a/b/c-OX*) lines, selected via neomycin (Neo)/phosphinothricin (PPT) resistance and validated by molecular genotyping and qPCR. For pot experiments, flowering Chinese cabbage seeds were germinated in darkness at 25°C for 12 h before sowing in sterilized substrate, with plants grown at 22-25°C, 70% humidity, 150 µmol m^-^²·s^-^¹ light intensity (16/8 h photoperiod), and watered with 0.1% nutrient solution every 3-5 days.^1^ For aseptic seedling growth assay, sterilized seeds were plated on 1/2 MS medium, 25°C, dark-germinated for 12 h, then grown for 2 days at 25°C in plant tissue culture laboratory prior to chemical treatment. All *A. thaliana* materials were in the Columbia-0 (Col-0) ecotype background. T-DNA insertion lines *atala3* (SALK_129494) were purchased from Arashare (https://www.arashare.cn/), Transgenic Col-0 lines expressing BraALA3a/b/c-GFP (*p35S::BraALA3a/b/c-GFP*) were selected by hygromycin (Hyg) resistance and PCR verification. For pot experiments, *A. thaliana* seeds were stratified at 4°C for 2 days before sowing on sterilized substrate, with plants grown at 20-22°C under otherwise identical conditions to those used for flowering Chinese cabbage. For aseptic seedling growth assay, surface-sterilized *A. thaliana* seeds were plated on 1/2 MS medium (1/2 MS, 3% sucrose, pH 5.8, 4 g/L agar), 4°C, darkness for 2 days, then grown at 22°C in plant tissue culture laboratory for 4 days before chemical treatment.

### PCZ Treatment and Phenotype Analysis

Flowering Chinese cabbage plants were treated with PCZ at bolting stage in pot experiments, with phenotype recorded post-treatment. PCA identified plant height and branch angle as representative metrics for genetic modification assessments. For aseptic seedling growth assay, 2-day-old seedlings were transferred to 1/2 MS medium containing PCZ, with hypocotyl and root elongation measured after 4 days.

Potted *A. thaliana* plants were treated with PCZ at 28 days post-sowing, with petiole length, leaf length, and leaf width measured in mature leaves post-treatment. For aseptic seedling growth assay, 4-day-old seedlings were transferred to 1/2 MS medium containing PCZ, and root elongation was quantified after 5 days of exposure.

### Measurement of PCZ Level by UPLC-MS/MS

The PCZ content was determined following the methodology described in previous literature.^7^ Detailed experimental procedures are provided in method S1 of the supporting information.

### Histological Section

Histological section were prepared following the methodology described in previous literature.^27^ Detailed experimental procedures are provided in method S2 of the supporting information.

### Analysis of Phylogenetic Tree

The DRS2 (YAL026C) sequence was searched on NCBI (https://www.ncbi.nlm.-nih.gov/) and blasted for homologous genes in Cruciferae species, multiple protein sequences were downloaded and phylogenetic tree analysis was performed using Poisson model based on neighbor-joining (NJ) method in MEGA 11.0 software to create Clusterw multiple sequence alignments.^28^

### Heterologous Expression in Yeast

Sensitivity testing of yeast with heterologous expression was performed following the methodology described in previous literature.^27,29,30^ Primers were shown in Table S2. Detailed experimental procedures are provided in method S3 of the supporting information.

### Generation of *BraALA3a/b/c^atala^*^3^ Complemented Lines

*A. thaliana* transformation was performed via floral dip using GV3101 harboring the 1300-BraALA3a/b/c-GFP construct.^31^ Detailed experimental procedures are provided in method S4 of the supporting information.

### Molecular Docking of BraALA3a/b/c-BraALIS2 with PCZ

The conformational structures of DRS2-Cdc50p protein were obtained from Protein Data Bank.^22^ ColabFold (https://colabfold.mmseqs.com/) was used for protein 3D structure modelling.^32^ The complex model was evaluated and optimised using the methods of Ramachandran.^33^ Based on previous reports,^22,34^ the parameters of the docking box were determined. Molecular docking was performed using Autodock Vina.^35,36^ Detailed experimental procedures are provided in method S5 of the supporting information.

### Molecular Dynamics Simulation (MD)

50 ns all-atom molecular dynamics simulations of the complex model were performed using the GROMACS software package.^37^ The stability of conformations was assessed by Root Mean Square Deviation (RMSD), Root Mean Square Fluctuation (RMSF), Radius of Gyration (Rg), Solvent-Accessible Surface Area (SASA) and number of hydrogen bonds.^38^ Detailed experimental procedures are provided in method S6 of the supporting information.

### P4-ATPase Protein Expression

In vitro protein expression experiments were conducted following the methodology described in previous literature.^39^ Detailed experimental procedures are provided in method S7 of the supporting information.

### Binding Affinity Determination by SPR

Protein-ligand interactions between BraALA3a/b/c-BraALIS2 dimers and PCZ were analyzed by SPR following the methodology described in previous literature.^40^ Detailed experimental procedures are provided in method S8 of the supporting information.

### *In Vitro* ATPase Activity Assay

Functional validation of protein complex bioactivity was performed through *in vitro* ATPase activity assays, including testing for potential P4-ATPase activators or inhibitors.^39^ Detailed experimental procedures are provided in method S9 of the supporting information.

### Quantitative Real-Time PCR (qRT-PCR)

RT-qPCR was performed following the methodology described in previous literature.^41^ *UBC10* served as the reference gene for flowering Chinese cabbage.^42^ The primers used for RT-qPCR analysis are listed in Table S2. Detailed experimental procedures are provided in method S10 of the supporting information.

### Generation of *BraALA3a/b/c*-Overexpressing Lines in flowering Chinese cabbage

*Agrobacterium*-mediated transformation of flowering Chinese cabbage was performed by cloning *BraALA3a/b/c* coding sequences into the pBI121 binary vector.^5^ Detailed experimental procedures are provided in method S11 of the supporting information.

### Generation of *BraALA3a/b/c* Knockdown Lines in flowering Chinese cabbage

*BraALA3a/b/c* knockdown lines in flowering Chinese cabbage were obtained following the methodology described in previous literature.^42^ Detailed experimental procedures are provided in method S12 of the supporting information.

### Physiological Parameter Quantification in flowering Chinese cabbage

Soluble protein content was determined by coomassie brilliant blue assay. Reduced vitamin C content was determined by molybdenum blue assay. Soluble sugar content was determined by anthrone assay. Chlorophyll content was determined by acetone/ethanol assay.^2,5^ All measurements included three biological replicates. Detailed experimental procedures are provided in method S13 of the supporting information.

### Establishment of a Rapid Screening System Based on Protoplast for Gene-Editing Targets in flowering Chinese cabbage

A rapid screening system based on protoplast for gene-editing targets in flowering Chinese cabbage was established following the methodology described in previous literature.^43,44^ Mutation profiles were characterized by Sanger sequencing or high-throughput sequencing,^45^ confirming editable targets. Detailed experimental procedures are provided in method S14 of the supporting information.

### Generation of *BraALA3a* Gene-Edited Lines in flowering Chinese cabbage

A CRISPR-Cas9 vector containing four tandem sgRNAs driven by the AtU6-26 promoter was constructed based on protoplast-validated editing targets, transformed into GV3101, and used to infect cotyledon-petiole explants of flowering Chinese cabbage. The transformation process involved explant preparation, pre-culture, infection, co-cultivation, antibacterial differentiation, antibiotic selection (MS, 3%Sucrose, 0.75%Agar, pH 5.8, 2 mg/L TZ, 0.5 mg/L IAA, 7 mg/L AgNO_3_, 6 mg/L Phosphinothricin [PPT]), shoot regeneration, root regeneration, and acclimatization.^5,46^ T0 plants were genotyped by high-throughput sequencing of pooled branch DNA, with editing profiles confirmed by Sanger sequencing. Plants were self-pollinated to propagate the progeny.

### Statistical Analysis

Data were presented as means ± standard deviations (SD) and calculated using triplicate measurements. Based on different comparisons, statistical significance was evaluated by two-way ANOVA (Dunnett/Šídák, \**P* < 0.05, \*\**P* < 0.01, \*\*\**P* < 0.001, no mark presents no significance), or one-way ANOVA (Tukey, *P* < 0.05). Statistical graphs were drawn using GraphPad Prism 10.3

## RESULTS

### PCZ Suppresses the Excessive Vegetative Growth of flowering Chinese cabbage

To identify the optimal application stage and concentration of PCZ for regulating excessive vegetative growth in flowering Chinese cabbage, we conducted a two-factor screening experiment. The results showed that plant height was significantly reduced when PCZ was applied at 25, 50, and 100 mg/L during the seedling (S1) stage, and at 50 and 100 mg/L during the bolting (S3) and harvest (S5) stages (Figure S1A). As the S3 stage marks the exponential phase of stem elongation,^47^ it was selected for further study. At this stage, both 50 and 80 mg/L PCZ induced significant dwarfing and stem thickening to a similar extent, yet residue levels were significantly lower at 50 mg/L (Figure S1B-D). Therefore, 50 mg/L PCZ was selected for subsequent pot experiments in flowering Chinese cabbage.

Principal component analysis (PCA) identified key agronomic traits affected by PCZ treatment, including plant height, internode length, main stalk stem length, and branch angles (Figure 1A, B). Accordingly, the representative growth parameters plant height and branch angle were selected for subsequent analysis. PCZ application induced pronounced dwarfism and architectural compactness, significantly reducing plant height and branch angles from 3 days after treatment onward (Figure 1C-E). Histological analysis of resin-embedded sections revealed that PCZ treatment compacted cell arrangement in the cortex, phloem, cambium, and xylem of the stem base (Figure S2A), increased cortical cell length while decreasing cell width (Figure S2B), and induced irregular deformation of medulla cells in petioles and leaves, accompanied by reduced cell length and width (Figure S2A, C, D). These alterations collectively demonstrate the efficacy of PCZ in suppressing excessive vegetative growth and shaping plant architecture in flowering Chinese cabbage.

**Figure 1.**
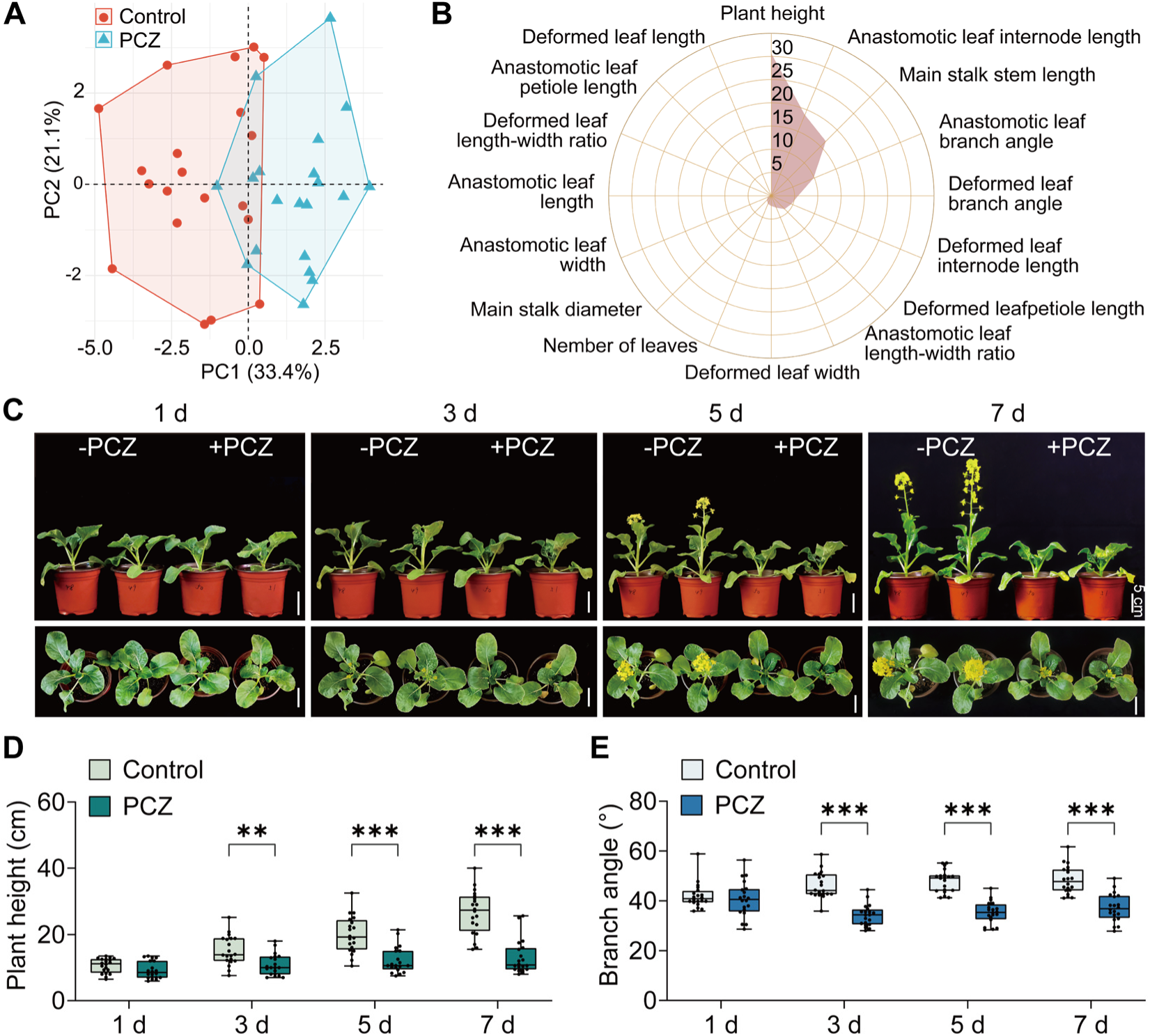
Growth inhibition of flowering Chinese cabbage by PCZ and principal component analysis (PCA) of growth indicators. (A) PCA of control-treated or 50 mg/L PCZ-treated flowering Chinese cabbage. (B) Phenotypic traits radar chart of control-treated or 50 mg/L PCZ-treated flowering Chinese cabbage. (C) 1, 3, 5, and 7 d phenotypes of control-treated or 50 mg/L PCZ-treated flowering Chinese cabbage. Bars = 5 cm. (D, E) Comparison of 1, 3, 5, and 7 d plant height (D) and branch angle (E) of control-treated or 50 mg/L PCZ-treated flowering Chinese cabbage. Data are mean ± SD (*n* = 20).

### Yeast Strains and *A. thaliana atala3* Mutant Heterologously Expressing the *BraALA3a/b/c* Gene Exhibited Sensitivity to PCZ

A previous screen of a membrane protein-deficient yeast library identified *drs2* mutant strains exhibiting resistance to PCZ. Phylogenetic and homologous comparative analyses identified three paralogous genes-*BraALA3a* (LOC103829275, Bra035422), *BraALA3b* (LOC103862629, Bra017897), and *BraALA3c* (LOC103843300, Bra031711) in flowering Chinese cabbage as homolog of yeast *DRS2* (YAL026C), sharing 44.14%, 44.15%, and 43.94% sequence similarity, respectively (Figure S3 and Table S1).

Heterologous expression of *BraALA3a/b/c* in both *drs2Δ* and BY4741 yeast strains conferred increased sensitivity to PCZ in a concentration-dependent manner. Yeast strains expressing *BraALA3a/b/c* exhibited significantly inhibited growth on PCZ-containing media compared to pYES2-empty vector controls, with overexpression strains showing more pronounced growth suppression (Figure 2A). Parallel liquid culture assays monitoring OD_600_ revealed identical trends, with *BraALA3*-expressing strains displaying markedly reduced proliferation rates versus three control groups (Figure 2B-G). These collective data demonstrate that *DRS2* deficiency confers PCZ resistance in yeast, heterologous expression of *BraALA3a/b/c* orthologs restores sensitivity to PCZ, *BraALA3a/b/c* overexpression enhances sensitivity.

**Figure 2.**
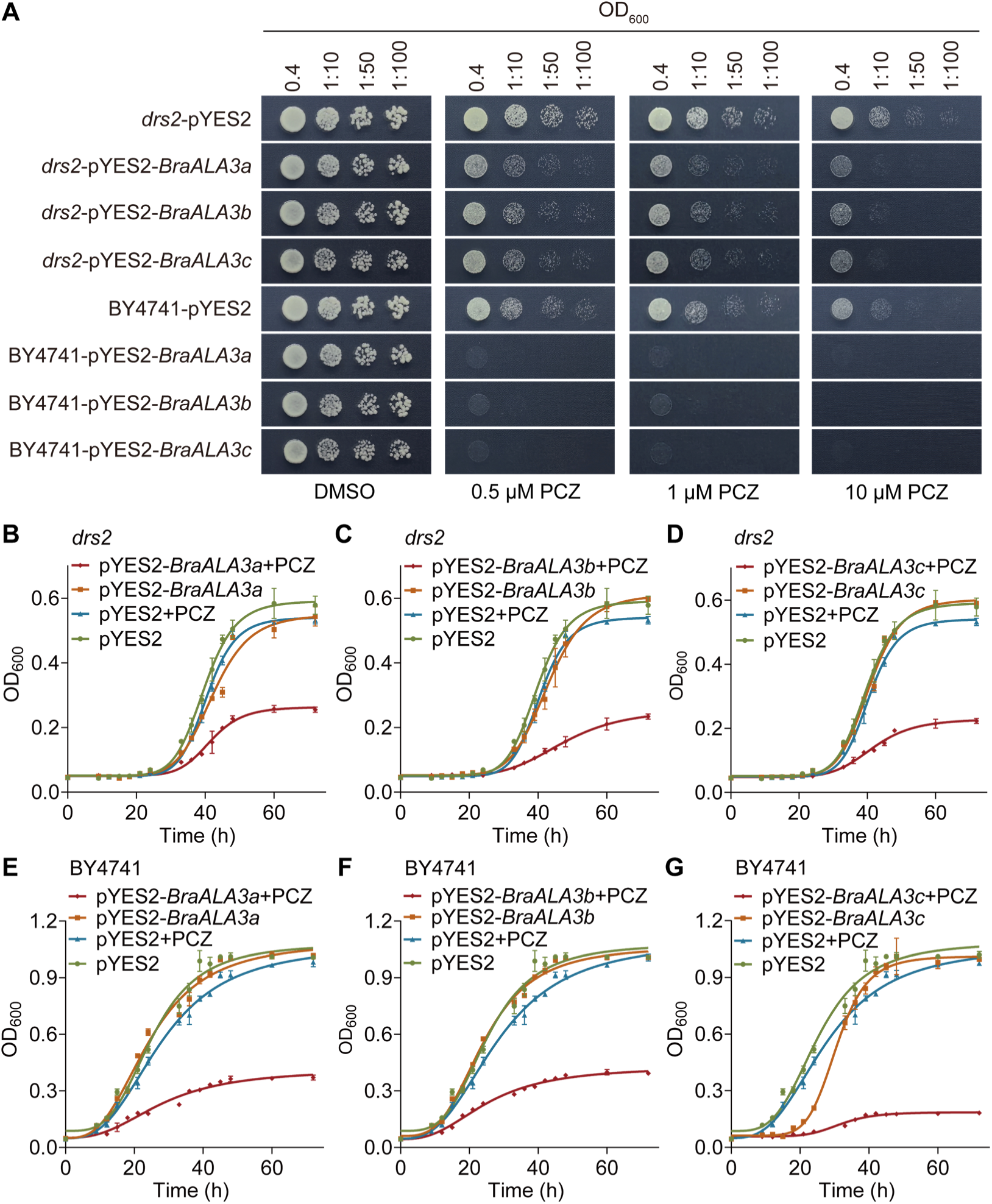
Identification of sensitivity of *BraALA3a/b/c* heterologous expression yeast strain to PCZ. (A) Growth of *drs2* and BY4741 yeast strains supplemented homologous gene *BraALA3a/b/c* on SD/-His-Leu-Met medium containing DMSO, 0.5 μM PCZ, 1.0 μM PCZ, or 10 μM PCZ. Plates were photographed after 3 days of incubation at 30°C. (B-D) Growth curve of *drs2* yeast strains supplemented homologous gene *BraALA3a/b/c* in SD/-His-Leu-Met liquid medium containing DMSO or 0.5 μM PCZ at 30°C. The growth of yeast strains was monitored by OD_600_ readings over 72 hours. Data are mean ± SD (*n* = 3). (E-G) Growth curve of BY4741 yeast strains supplemented homologous gene *BraALA3a/b/c* in SD/-His-Leu-Met liquid medium containing DMSO or 0.5 μM PCZ at 30°C. The growth of yeast strains was monitored by OD_600_ readings over 72 hours. Data are mean ± SD (*n* = 3).

Heterologous expression of *BraALA3a/b/c* in *A. thaliana atala3* mutants and subsequent phenotypic revealed that complementation lines rescued the growth defects in both seedlings and mature plants (Figure S4A, B). Compared to Col-0, PCZ-treated *atala3* plants showed reduced suppression of petiole length, leaf length and width (Figure S4C-E), confirming *AtALA3* deficiency decreases PCZ sensitivity. Complemented lines expressing *BraALA3a/b/c* exhibited stronger PCZ-induced suppression of these parameters than *atala3* (Figure S4C-E), indicating that the expression of *BraALA3a/b/c* restored sensitivity to PCZ, demonstrating functional conservation of PCZ responsiveness between orthologs. Similarly, root length analysis in seedlings demonstrated the same trend (Figure S4F), which is consistent with the sensitivity to PCZ observed in the heterologously expressing yeast (Figure 2).

### PCZ Competes with PI4P for Binding to the Active Pocket of BraALA3a/b/c Proteins

The functional activity of P4-ATPases is dependent on their interaction with a *β*-subunit.^20,21^ Using the DRS2-Cdc50p structure as a template,^22^ homology models of BraALA3a/b/c-BraALIS2 complexes were generated. Molecular docking results of BraALA3a/b/c-BraALIS2 with PCZ revealed that PCZ binds within a hydrophobic pocket formed by transmembrane helices TM7, TM8, and TM10, along with the autoinhibitory C-terminus, establishing an interaction conformation (Figure 3A-C). The BraALA3a-BraALIS2 complex exhibited a binding energy of -8.2 kcal/mol with PCZ, corresponding to a ligand efficiency of 0.3737 kcal/(mol·K). The triazole ring of PCZ forms hydrogen bonds with Asparagine 1045 (Asn-1045) and Glutamine 1129 (Gln-1129), while also engaging in both a hydrogen bond and a π-π interaction with Tyrosine 1125 (Tyr-1125). The dioxolane group forms hydrophobic-interaction with Leucine 1042 (Leu-1042), Aspartic acid 1110 (Asp-1110), and Gln-1114. Additionally, the para-substituted phenyl ring and propyl side chain form halogen bond with Gln-1114 and Proline 1253 (Pro-1253), as well as a hydrophobic-interaction with Leu-1255 (Figure 3A and Figure S5A-C). The BraALA3b-BraALIS2 complex exhibited a binding energy of -7.1 kcal/mol with PCZ, corresponding to a ligand efficiency of 0.3227 kcal/(mol·K).

**Figure 3.**
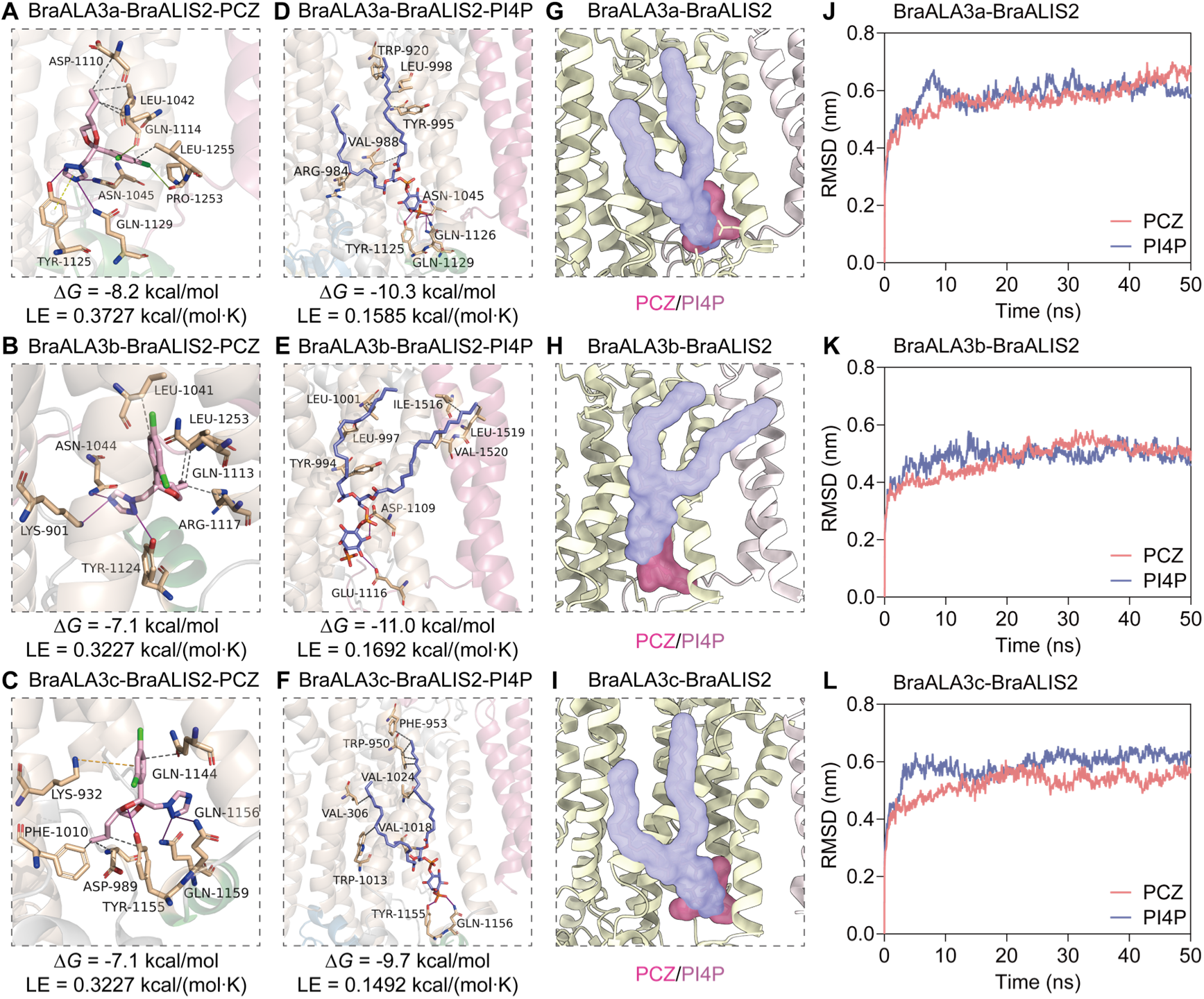
Molecular docking of BraALA3a/b/c-BraALIS2 protein binding to PCZ/PI4P. (A-C) Binding site visualization of molecular docking between BraALA3a/b/c-BraALIS2 and PCZ. (D-F) Binding site visualization of molecular docking between BraALA3a/b/c-BraALIS2 and PI4P. △*G* represents binding energy. LE represents ligand efficiency. PCZ (pink) and PI4P (purple) are represented by the stick model. Gray dashed lines represent hydrophobic-Interactions, purple solid lines represent hydrogen-bonds, yellow dashed lines represent π-stacking, green solid lines represent halogen-bonds, orange dashed lines represent π-cation-interactions, yellow solid lines represent salt-bridges. (G-I) Schematic representation of spatial structure of PCZ (pink) and PI4P (purple) robbing BraALA3a/b/c-BraALIS2 pockets. (J-L) The Root Mean Square Deviation (RMSD) curves of BraALA3a/b/c-BraALIS2 and PCZ/PI4P complexes over time.

The N atom of the triazole ring in PCZ forms hydrogen bonds with Asn-1044, Lysine 901 (Lys-901), and Tyr-1124. The dioxolane group forms hydrophobic-interactions with Gln-1113 and Arginine 1127 (Arg-1127), as well as Leu-1253. Additionally, the para-substituted phenyl ring and propyl side chain forms hydrophobic-interactions with Leu-1041 (Figure 3B and Figure S5A-C). The BraALA3c-BraALIS2 complex displayed a binding energy of -7.1 kcal/mol with PCZ, corresponding to a ligand efficiency of 0.3227 kcal/(mol·K). The N atom of PCZ’s triazole ring forms hydrogen bonds with Gln-1156 and Gln-1159, while its dioxolane group forms hydrophobic-interactions with Asp-989, Phenylalanine 1010 (Phe-1010), and Tyr-1155. The para-substituted phenyl ring and propyl side chain form a π-cation interaction with Lys-932 and a hydrophobic interaction with Gln-1144 (Figure 3C and Figure S5A-C). These results indicate that the triazole ring of PCZ primarily interacts with BraALA3a/b/c via hydrogen bonds, whereas the dioxolane moiety predominantly forms hydrophobic-interactions with BraALA3a/b/c.

To function, P4-ATPases must be activated in a PI4P-dependent manner, this activation enables the flipping of specific phospholipid substrates such as PS across the membrane.^22,48^ Homology modeling and molecular docking revealed that both PI4P and PCZ bind to the same active site in BraALA3a/b/c, formed by transmembrane helices TM7, TM8, TM10, and the autoinhibitory C-terminal pocket (Figure 3D-F and Figure S5D-F). Co-docking confirmed shared occupation of this pocket (Figure 3G-I and Figure S5), with key common interacting residues identified (e.g., Asn-1045, Tyr-1125, Gln-1129 in BraALA3a; Tyr-1155, Gln-1156 in BraALA3c). Although the binding affinity of PCZ to BraALA3a/b/c was marginally lower than that of PI4P, its ligand efficiency significantly surpassed that of PI4P (Figure 3A-F). Molecular dynamics simulations demonstrated stable complex formation and comparable dynamic trajectories for both ligands (Figure 3J-L and Figure S6). Free energy decomposition highlighted residues with major binding energy contributions (≤ –10 kcal/mol) for both PCZ and PI4P across the three homologs (Figure S7). Notably, several residues contributed more strongly to PCZ binding (Figure S7), supporting a model in which PCZ competitively occupies the PI4P-binding pocket through non-covalent interactions at key sites within the active pocket.

### PCZ Directly Interacts with BraALA3a/b/c Proteins and Inhibits Their ATPase Activity

To validate the interaction predicted by molecular docking, the BraALA3a/b/c proteins were expressed in the BJ5465 yeast system, and their binding affinity for PCZ was determined using surface plasmon resonance (SPR). SDS-PAGE confirmed successful expression of BraALA3a/b/c membrane proteins in flowering Chinese cabbage (Figure S8A). Compared with the control DMSO (Figure S8B-D), concentration-dependent SPR binding signals (10 nM to 2560 nM) demonstrated enhanced PCZ interactions with BraALA3a/b/c homologous proteins, generating characteristic SPR binding kinetics curves (Figure 4A-C). The equilibrium dissociation constants (Avg KD) for BraALA3a and BraALA3c were 6.14 × 10^-8^ M and 1.40 × 10^-7^ M respectively, indicating strong binding affinities, while BraALA3b showed middle binding with an Avg KD of 1.05 × 10^-5^ M (Table S3). These results demonstrate significant binding affinities between ALA3 membrane proteins and PCZ, with BraALA3a exhibiting the strongest interaction in flowering Chinese cabbage, followed by BraALA3c, while BraALA3b showed relatively weaker binding.

**Figure 4.**
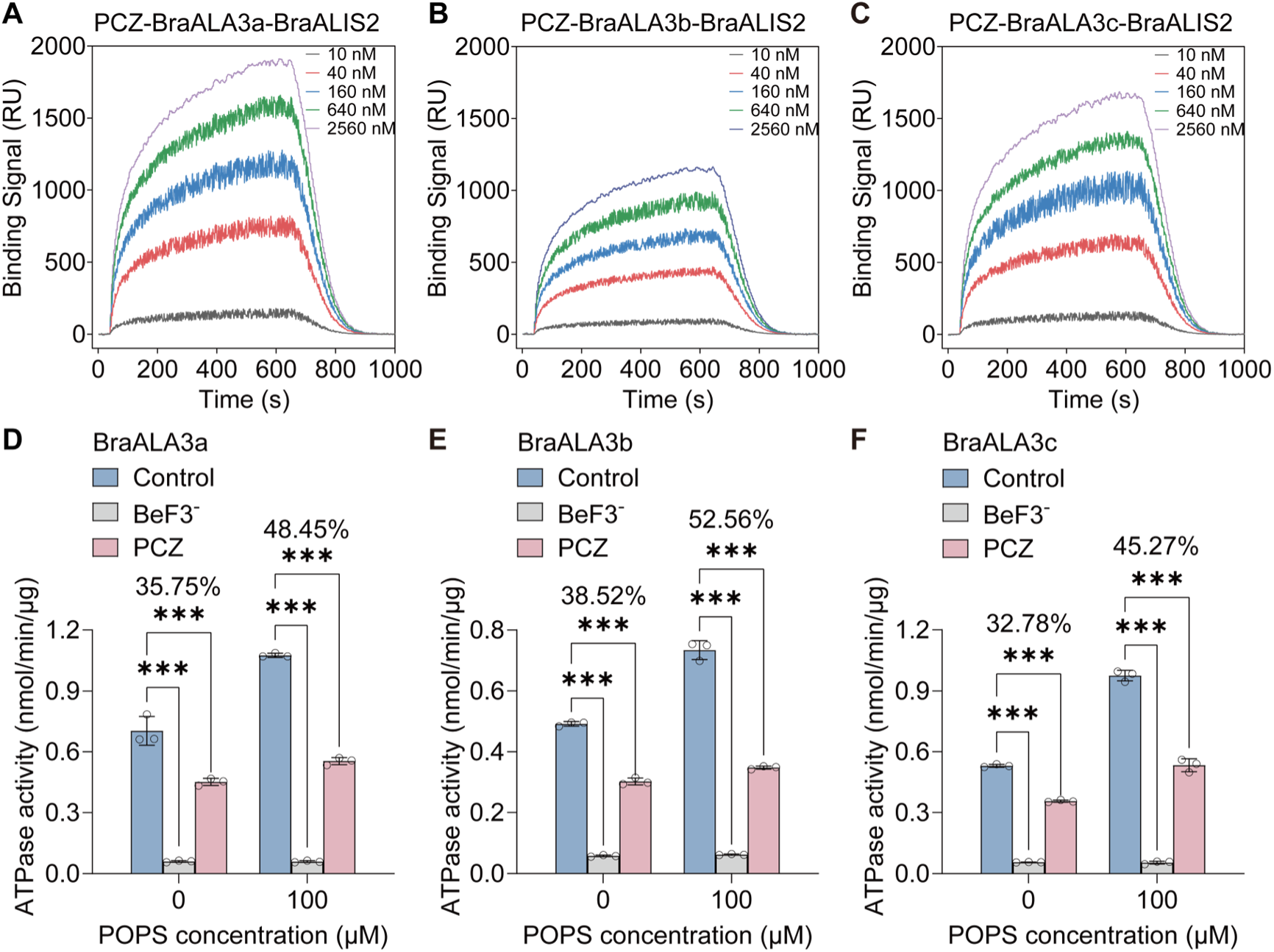
Validation of PCZ-BraALA3a/b/c binding affinity and PCZ-mediated ATPase activity inhibition. (A-C) Surface plasmon resonance (SPR) affinity binding signals of BraALA3a/b/c-BraALIS2 and PCZ. (D-F) Functional validation of P4-ATPase protein complex enzymatic activity. BeF^-^ acts as an inhibitor of P-type ATPases. POPS is phosphatidylserine. Data are mean ± SD (*n* = 3). Significance differences were tested by two-way ANOVA (Dunnett, \**P* < 0.05, \*\**P* < 0.01, \*\*\**P* < 0.001).

To determine whether PCZ acts as an inhibitor or activator of BraALA3a/b/c, *in vitro* ATPase activity assays was performed. As expected, the known P-type ATPase inhibitor BeF₃⁻ served as an effective negative control, reducing ATPase activity to basal levels (Figure 4D-F). Compared to the untreated control, PCZ significantly inhibited the ATPase activity of BraALA3a, BraALA3b, and BraALA3c by 35.75%, 38.52%, and 32.78%, respectively (Figure 4D-F). Since the substrate PS is known to stimulate flippase activity,^39,49^ its effect was assessed. The addition of PS markedly enhanced the ATPase activity of BraALA3a/b/c homologs. Under this activated condition, the inhibitory effect of PCZ was substantially potentiated, with suppression rates increasing to 48.45%, 52.56%, and 45.27% for BraALA3a/b/c, respectively (Figure 4D-F). These results demonstrate that PCZ is an inhibitor of BraALA3a/b/c ATPase activity, and its efficacy is enhanced in the presence of the lipid substrate PS.

### PCZ Affects Transcriptional Regulation Patterns of *BraALA3a/b/c*Genes

To assess the effect of PCZ on transcript levels, qRT-PCR was performed. Compared to untreated controls, the expression of *BraALA3a* and *BraALA3b* peaked at 24 h, while *BraALA3c* transcript levels remained elevated at 72 h in roots (Figure S9A-C). *BraALA3a* and *BraALA3c* showed sustained high expression at post-treatment while *BraALA3b* displayed transient induction peaking at 12 h before returning to baseline in stems (Figure S9D-F). Mature leaves exhibited acute transcriptional activation of *BraALA3a/b/c* orthologs at 3 h post-treatment (Figure S9G-I), whereas new leaves showed a biphasic response characterized by initial suppression at 24 h followed by rebound overexpression (Figure S9J-L). These results indicate that PCZ affects the transcriptional expression of *BraALA3a/b/c*, with a more persistent impact in newly developed tissues like young leaves. This suggests that the PCZ-responsive expression of *BraALA3a/b/c* genes may be an important factor influencing the growth of nascent tissues.

### Overexpression of *BraALA3a/b/c* Enhanced Sensitivity to PCZ in flowering Chinese cabbage, Whereas Knockdown of These Genes Reduced Sensitivity to PCZ

To assess the phenotypic sensitivity of flowering Chinese cabbage to PCZ, *Agrobacterium*-mediated overexpression system was established in flowering Chinese cabbage by transforming explants with pBI121 vectors carrying target genes. *BraALA3a/b/c* overexpression lines were identified by PCR (Figure S10A), qRT-PCR identification results showed that the upregulation of *BraALA3a* transcription expression was up to 14.30-fold, *BraALA3b* expression was up to 17.12-fold, and *BraALA3c* expression was up to 11.59-fold, while revealing coordinated upregulation among paralogs (Figure S10B-D). *OXBraALA3a21-3*, *OXBraALA3b3-2*, and *OXBraALA3c1-4* were propagated to T2 generation. Phenotypic analysis demonstrated *BraALA3a/b/c-OX* plants exhibited enhanced growth (Figure 5A), including increased plant height (Figure 5B) and wider branch angles (Figure 5C). Furthermore, these plants displayed thinner main stalk diameters (Figure S11A) and longer deformed leaf petioles (Figure S11B), whereas leaf dimensions remained largely unaffected (Figure S11C, D). PCZ treatment caused greater suppression of height and branch angle expansion in *BraALA3a-OX* compared to WT (Figure 5B, C), indicating heightened PCZ sensitivity-particularly in *BraALA3a-OX*.

**Figure 5.**
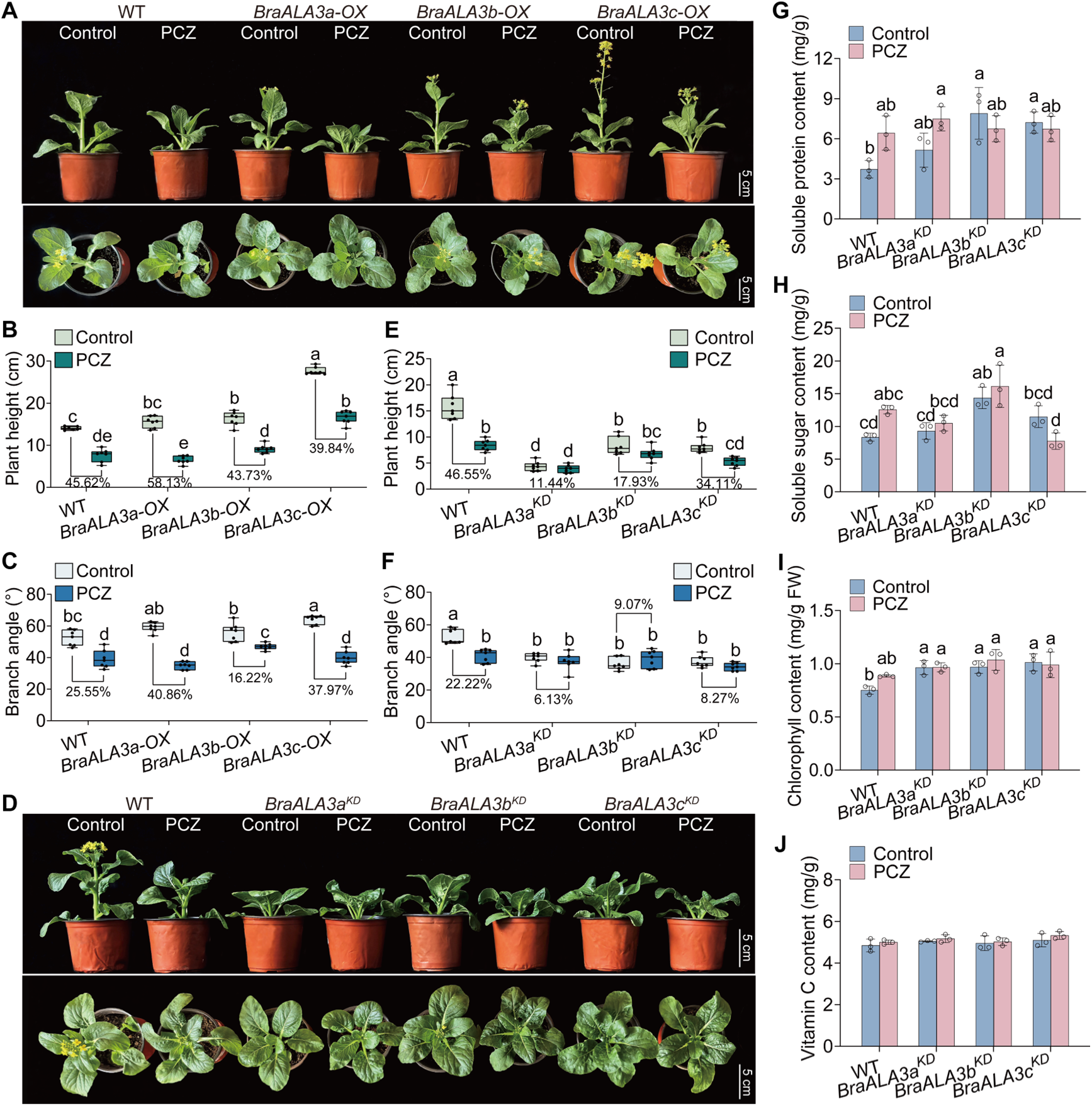
Phenotypic analysis of sensitivity to PCZ in *BraALA3a/b/c-OX* and *BraALA3a/b/c^KD^*. (A) 5 d phenotypes of control-treated or 50 mg/L PCZ-treated WT and *BraALA3a/b/c-OX*. Bars = 5 cm. *BraALA3a/b/c-OX* represents gene overexpressed flowering Chinese cabbage. (B, C) Comparison of 5 d plant height (B) and branch angle (C) of control-treated or 50 mg/L PCZ-treated WT and *BraALA3a/b/c-OX*. Data are mean ± SD (*n* = 7). (D) 5 d phenotypes of control-treated or 50 mg/L PCZ-treated WT and *BraALA3a/b/c^KD^*. Bars = 5 cm. *BraALA3a/b/c^KD^*represents gene knocked down flowering Chinese cabbage. (E, F) Comparison of 5 d plant height (E) and branch angle (F) of control-treated or 50 mg/L PCZ-treated WT and *BraALA3a/b/c^KD^*. Data are mean ± SD (*n* = 7). (G-J) Comparison of 5 d soluble protein content (G), soluble sugar content (H), chlorophyll content (I), and reduced vitamin C content (J) of control-treated or 50 mg/L PCZ-treated WT and *BraALA3a/b/c^KD^*. Significance differences were tested by one-way ANOVA (Tukey). Different letters represent significant differences (*P* < 0.05). Percentages represent the decrease or increase of each indicator.

*Agrobacterium*-mediated gene knockdown system was established in flowering Chinese cabbage, constructing pBI121-amiRNA vectors (Figure S12A) for transcriptional silencing of *BraALA3a/b/c* via vacuum-infiltration floral dip. Transgenic lines were verified by PCR (Figure S12B), with qRT-PCR revealing compensatory expression between *BraALA3a/BraALA3b* and coordinated regulation between *BraALA3a/BraALA3c* (Figure S12C-E). T1 seeds were collected from selected lines (*BraALA3a^KD^*-13, *BraALA3b^KD^*-6, *BraALA3c^KD^*-5). Phenotypic analysis showed *BraALA3a/b/c* knockdown lines exhibited growth inhibition (Figure 5D) with reduced plant height (Figure 5E) and narrower branch angles (Figure 5F). Furthermore, these plants displayed thicker main stems (Figure S13A), whereas petiole length, leaf length, and leaf width showed no significant alterations (Figure S13B-D). Compared to WT, PCZ-treated *BraALA3a/b/c^KD^*plants showed less suppression of height and branch angle (Figure 5E, F), indicating reduced sensitivity to PCZ - particularly in *BraALA3a^KD^*. These findings are consistent with both the phenotypic observations from *BraALA3a/b/c* overexpression plants (Figure 5A-C) and the higher binding affinity of BraALA3a for PCZ as determined by SPR (Figure 4A-C and Table S3). Physiological assays revealed that the *BraALA3a/b/c* knockdown lines exhibited increased contents of soluble proteins, soluble sugars, and chlorophyll, while the level of Vitamin C remained unaltered (Figure 5G-J). Collectively, these results demonstrate that down-regulating *BraALA3a/b/c* not only effectively restrains excessive vegetative growth but also maintains or even enhances key nutritional components.

In conclusion, phenotypic analyses in flowering Chinese cabbage demonstrate that overexpression of *BraALA3a/b/c* enhances sensitivity to PCZ (Figure 5A-C), while knockdown of these genes reduces sensitivity to PCZ (Figure 5D-F). Furthermore, among its homologs, *BraALA3a* confers the highest responsiveness to PCZ (Figure 5), it was selected for subsequent gene editing.

### Heterozygous Gene-Edited Lines of *BraALA3a* in flowering Chinese cabbage Was Generated

To establish *BraALA3a* gene-edited plants, a rapid target screening system based on protoplast in flowering Chinese cabbage was first developed, comprising: *in vitro* sgRNA transcription and cleavage assays, RNP transfection into protoplasts, and editing efficiency validation of RNP induced mutations. Protoplasts were isolated from flowering Chinese cabbage using an enzymatic digestion method (Figure 6A), protoplast viability was confirmed by FDA staining (Figure 6B), with transfection competence verified by GFP expression (Figure 6C). Post-RNP transfection, cells cultured in 1N0.3K medium maintained division and differentiation capacity (Figure 6D). PCR and sequencing of proliferated cells identified editing targets, while *BraA06·GA4* editing efficiency and double-strand break (DSB) site (Figure S14A) matched published patterns,^43^ validating our platform. Three editable active sites within the *BraALA3a* gene were identified, corresponding to sgRNA02 (Figure S14B), sgRNA03 (Figure 6E), and sgRNA07 (Figure S14C).

**Figure 6.**
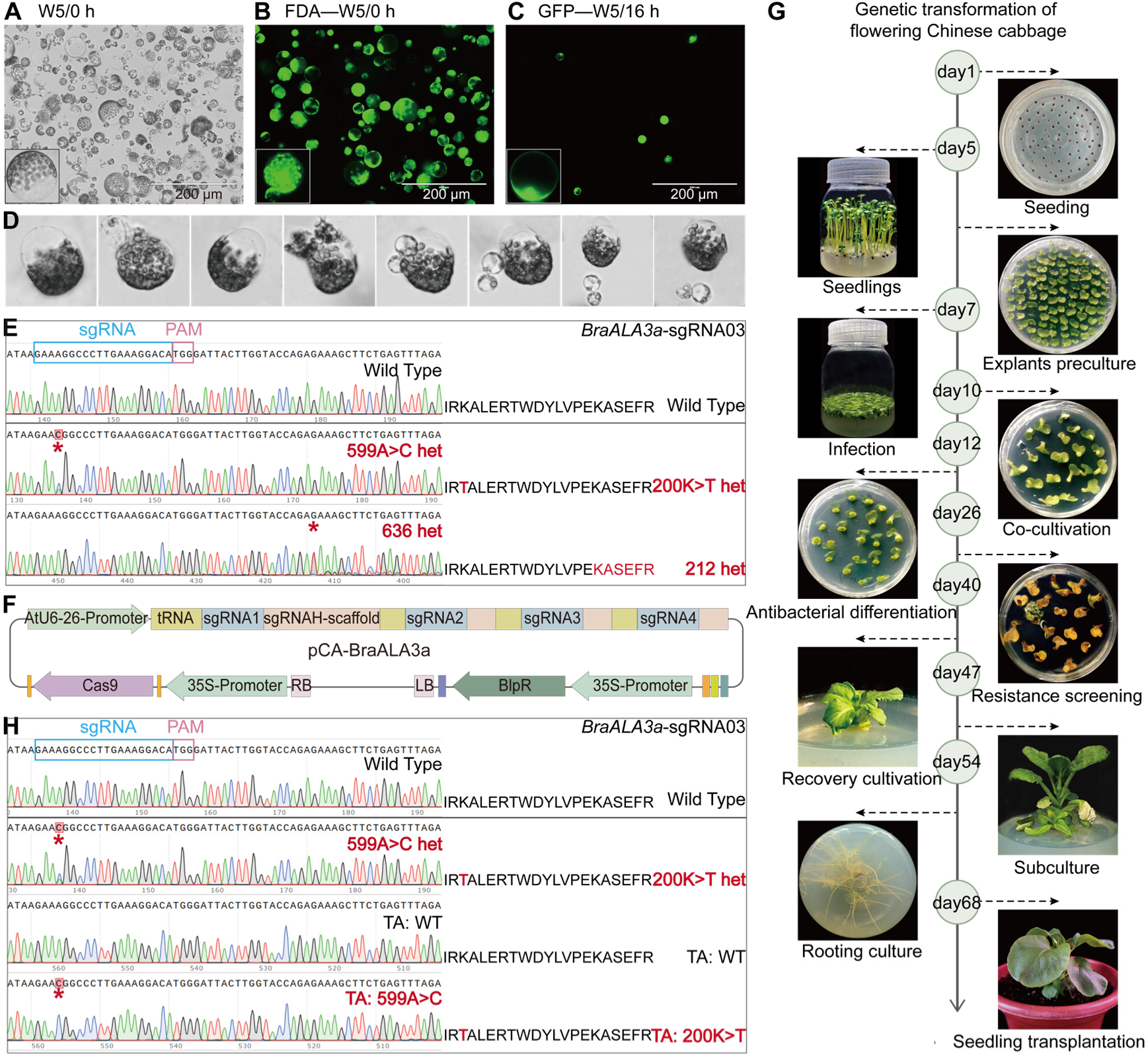
CRISPR-Cas9 genome editing via protoplast-based and *Agrobacterium tumefaciens*-mediated in flowering Chinese cabbage. (A) Freshly isolated protoplasts. Bar = 200 µm. (B) Viable protoplasts identified by Fluorescein diacetate (FDA) staining. Bar = 200 µm. (C) Transfected protoplasts expressing GFP protein observed under fluorescence microscope. Bar = 200 µm. (D) Protoplasts undergoing cell divisions and multiplication. (E) Sanger sequencing verification of target site editing profiles in protoplast. (F) Architecture of CRISPR-Cas9 editing vector targeting *BraALA3a* in flowering Chinese cabbage. (G) Schematic of *Agrobacterium tumefaciens*-mediated genetic transformation in flowering Chinese cabbage. (H) Sanger sequencing verification of target site editing profiles in explant. PAM sites are highlighted in pink, sgRNA are highlighted in blue, mutated nucleotides were highlighted in red.

Using a CRISPR/Cas9 vector with four tandem sgRNAs (Figure 6F), *BraALA3a*-edited flowering Chinese cabbage was generated via *Agrobacterium*-mediated transformation of explants (Figure 6G). The resulting mutants included one carrying an A>C substitution at nucleotide position 599, resulting in a lysine to threonine substitution at amino acid 200 (Figure 6H), and the other exhibiting double peaks in the sequencing chromatogram starting from nucleotide 636 (Figure S15). Unfortunately, no homozygous lines were obtained through selfing in subsequent breeding generations, with only the BraALA3a/*braala3a^K200T^* heterozygous line remaining viable, which exhibited smaller seeds (Figure 7A), reduced germination rates (Figure 7B), and shorter post-germination length of root and bud (Figure 7C) compared to WT. The BraALA3a/*braala3a^K200T^* mutant exhibited a profound dwarfism from bolting through harvest (Figure 7D-F), characterized by significantly decreased plant height (Figure 7G), internode length (Figure 7H), and branch angle (Figure 7I), which collectively conferred strongly suppressed bolting and compact architecture. This consistent suppression of growth across developmental stages underscores the critical role of *BraALA3a* in promoting vegetative growth.

**Figure 7.**
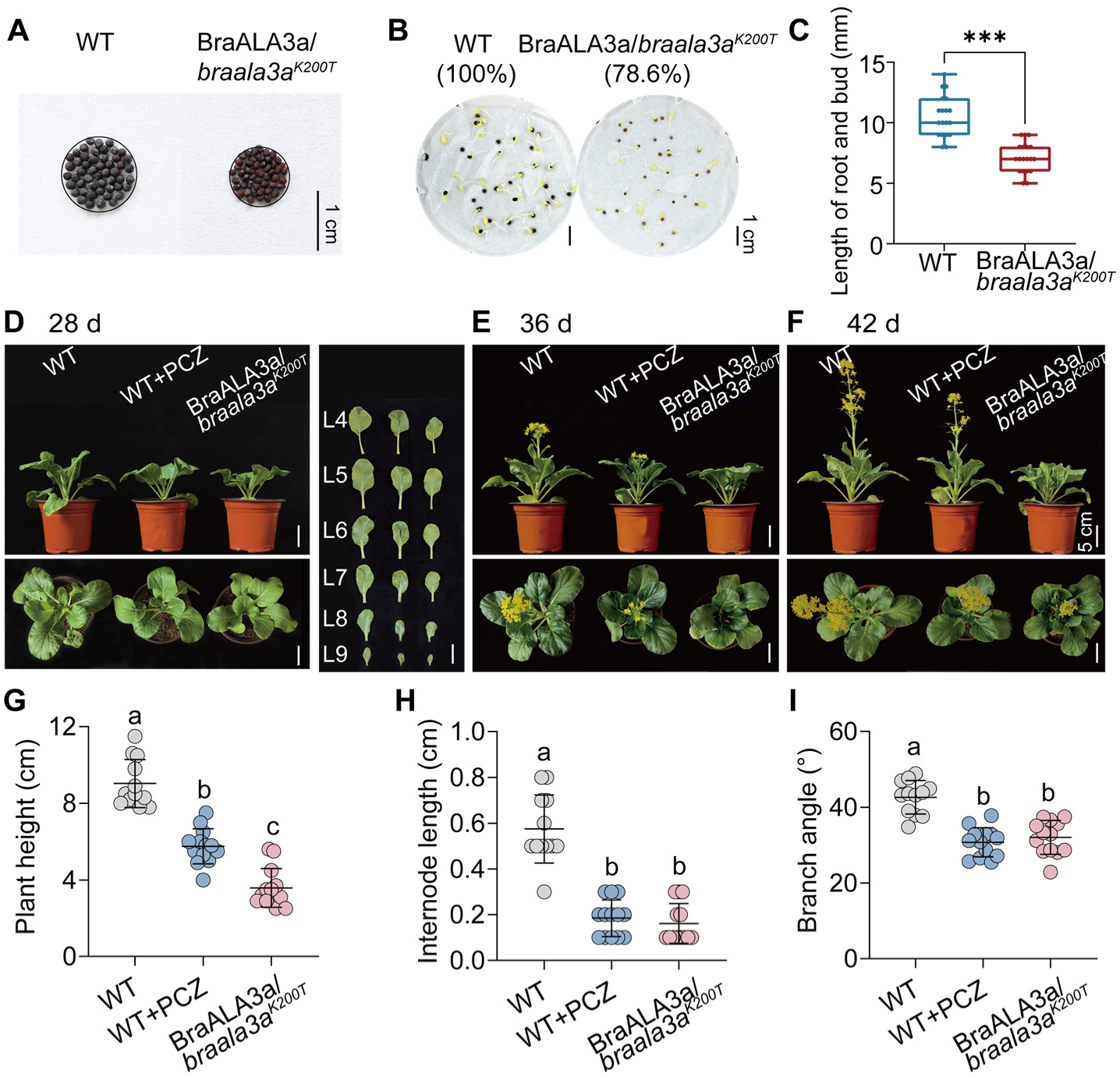
Phenotypes of BraALA3a/*braala3a^K200T^*. (A) Seed size of WT and BraALA3a/*braala3a^K200T^*mutant. (B) Germination rate of WT and BraALA3a/*braala3a^K200T^* mutant. (C) Root and shoot elongation of WT and BraALA3a/*braala3a^K200T^* mutant seeds at 48 hours post-germination. Data are mean ± SD (*n* = 19). Significance differences were tested by t-test (\*\*\**P* < 0.001). (D-F) 28 d (D), 36 d (E), and 42 d (F) phenotypes of WT and BraALA3a/*braala3a^K200T^* mutant. L4-L9 represent the newly formed leaves after the fourth leaf (i.e. deformed leaf) during the growth process of flowering Chinese cabbage. Bars = 5 cm. (G-I) Comparison of 28 d plant height (G), internode length (H), and branch angle (I) of WT and BraALA3a/*braala3a^K200T^* mutant. Data are mean ± SD (*n* ≥ 12). Significance differences were tested by one-way ANOVA (Tukey). Different letters in the same group data represent significant differences (*P* < 0.05).

## DISCUSSION

Minimizing irrational pesticide use remains a central challenge in modern agriculture, with profound significance for environmental conservation and the reduction of potential health risks. In flowering Chinese cabbage production, growers apply PCZ to optimize plant architecture by controlling excessive growth.^7,8^ However, the brief growth period of flowering Chinese cabbage imposes a significant constraint on the safe preharvest interval for PCZ, resulting in pesticide accumulation and residues that threatening dietary safety.^50,51^ Elucidating the molecular mechanism by which PCZ regulates plant architecture and establishing a safe, sustainable growth-control strategy will provide a critical scientific foundation for reducing and replacing PCZ reliance in agricultural practice. Triazole fungicides are known to inhibit fungal CYP51 (14α-demethylase), thereby blocking ergosterol biosynthesis and compromising membrane integrity to achieve antifungal effects.^11,12^ In plants, PCZ reportedly inhibits C22 and C23 side-chain hydroxylation during the conversion of campesterol to teasterone.^14^ However, the mechanism by which PCZ, as an exogenous compound, interacts with membrane structures upon cellular entry remains unclear, precluding a definitive understanding of its mode of action. A key breakthrough of this study is the identification of ALA3, a member of the P4-ATPase family, as a novel membrane-bound target of PCZ in plants. This finding is robustly supported by multi-layered evidence underscoring the functional significance of ALA3.

Phospholipid flippases of P4-ATPases in eukaryotes, catalyzing the translocation of phospholipids, such as phosphatidylserine and phosphatidylethanolamine, from the exoplasmic to the cytoplasmic leaflet of the lipid bilayer. Their activity is critical for maintaining membrane asymmetry and facilitating vesicle formation and trafficking.^22,48^ The ATPase activity of these flippases depends on the lipid regulator PI4P,^48^ which binds to a hydrophobic pocket formed by transmembrane helices TM7, TM8, and TM10 in the yeast P4-ATPase Cdc50p complex.^22^ The yeast DRS2 homologs in flowering Chinese cabbage are the membrane-localized BraALA3a/b/c proteins (Figure S3). PCZ exhibits nanomolar-scale binding affinity for BraALA3a/b/c (Figure 4A-C and Table S3), engaging a hydrophobic pocket formed by transmembrane helices TM7, TM8, TM10, and the autoinhibitory C-terminus (Figure 3A-C and Figure S5A-C). Notably, the lipid flippase regulator PI4P binds to an overlapping site within the same pocket (Figure 3D-F and Figure S5D-F). Thus, we propose that PCZ impedes the activation of the protein’s auto-regulatory conformational mechanism by competing with PI4P for binding to this transmembrane domain pocket (Figure 3G-I). The role of inhibitor is well-established in both basic research and agricultural practice. For instance, the compound 11e might bind to the BP2 pocket of *Xoo*FtsZ, inducing conformational changes that inhibit its GTPase activity and disrupted protofilament assembly, ultimately leading to bacterial death.^38^ In line with this, our *in vitro* assays demonstrated that PCZ suppresses the ATPase activity of BraALA3a/b/c proteins (Figure 4D-F), corroborating structural evidence suggesting that PCZ acts as a competitive inhibitor of ALA3 (Figure 8). Competitive inhibition leading to the loss of ALA3 function is expected to disrupt membrane lipid asymmetry, thereby directly impairing vesicular trafficking processes that depend on specific lipid environments.^25^ These findings provide a mechanistic explanation for the rapid PCZ-induced alterations in membrane architecture and fill a critical gap in understanding its mode of action from extracellular entry to intracellular.

**Figure 8.**
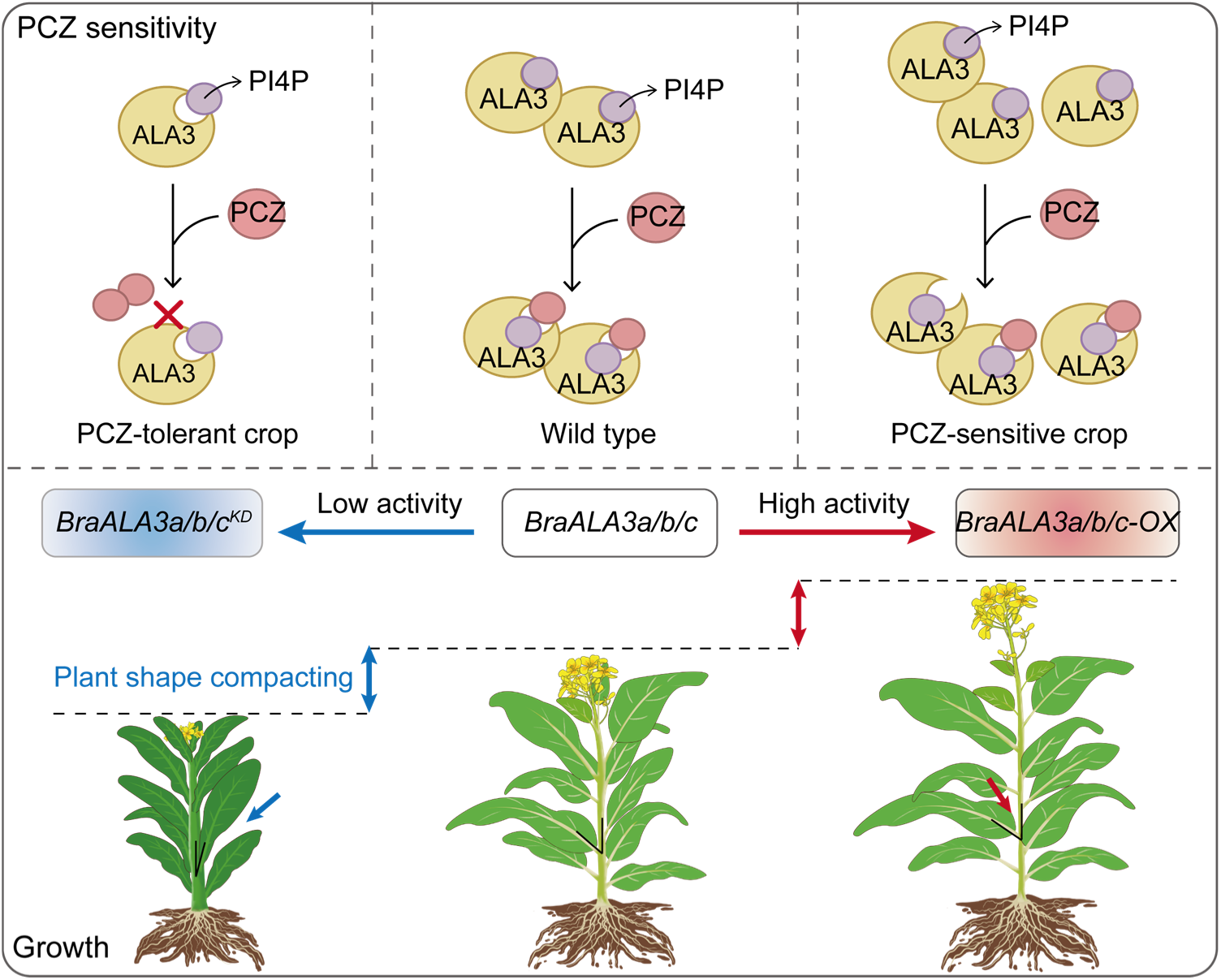
Mechanism model of PCZ targeting BraALA3a/b/c to regulate the growth of flowering Chinese cabbage. In the schematic, the yellow crescent represents the ALA3 protein; the red circle denotes the PCZ pesticide molecule; and the purple circle indicates the PI4P substrate molecule. Blue arrows (in different positions) signify downregulation of gene expression, reduced plant height, or smaller branch angles. Red arrows (in different positions) denote upregulation of gene expression, increased plant height, or larger branch angles.

Building on the mechanistic foundation of PCZ-BraALA3a/b/c interaction, in planta evidence further demonstrates that PCZ induces a rapid, stress-responsive upregulation of *BraALA3a/b/c* transcription (Figure S9). At the genetic level, overexpression of *BraALA3a/b/c* promoted plant growth and enhanced sensitivity to PCZ, whereas knockdown resulted in dwarfism and reduced sensitivity to PCZ (Figure 5). Based on these findings, we pursued the development of compact, architecture-optimized varieties of flowering Chinese cabbage through gene editing of *BraALA3a/b/c*, aiming to achieve green growth regulation without the need for PCZ application. The application of gene editing in flowering Chinese cabbage has rarely been reported. In one study, further optimization by integrating sgRNA and Cas9 expression cassettes from the psgR-Cas9-At system into the pBI121 vector achieved 20-50% editing efficiency in flowering Chinese cabbage. This system also facilitated large-fragment deletions in *BcPME37c* and *BcPME37b* and simultaneous editing of both genes.^52^ In another study, the CRISPR/Cas9 system was employed to generate efficient and heritable mutations in the key DELLA protein gene *BraRGL1* in flowering Chinese cabbage, yielding loss-of-function variants with two amino acid substitutions in the GRAS domain.^5^ In a novel approach, a method was developed to isolate, transfect, and regenerate individual mutagenized protoplasts into mature plants. This system used PEG-mediated delivery of Cas9/sgRNA constructs or ribonucleoprotein (RNP) complexes into *Brassica oleracea* protoplasts, achieving a targeted insertion frequency of 13.6% for the *PDS*, *GA4*, and *SRK* genes.^43,44^

Using a screening system involving *in vitro* transcription and cleavage assays of sgRNAs, RNP assembly, and transfection of flowering Chinese cabbage protoplasts, mutable target sites in the *BraA06·GA4* gene and three editable active sites in the *BraALA3a* gene were identified (Figure 6A-E and Figure S14). Stable genetic transformation via *Agrobacterium* tumefaciens was then performed, generating T0 heterozygous plants carrying an A>C substitution at nucleotide position 599 in the *BraALA3a* coding region, resulting in a lysine-to-threonine amino acid change at residue 200 (Figure 6F-H). The BraALA3a/*braala3a^K200T^*mutants exhibited a more dwarfed stature, significantly suppressed bolting, and compact plant architecture. Growth inhibition was observed consistently from sprouting to seedling establishment (Figure 7). These observations indicate that *BraALA3a* plays a critical role in regulating growth and development in flowering Chinese cabbage. Even a single amino acid substitution (K200T) is sufficient to induce multi-faceted growth inhibition phenotypes, suggesting that the gene likely influences fundamental cellular processes, such as membrane lipid asymmetry and vesicular trafficking, which in turn modulates the signaling and distribution of phytohormones including brassinosteroids and auxins, ultimately leading to altered plant architecture and developmental timing. Alternatively, the *BraALA3a* mutation may represent a dominant-negative or gain-of-function allele. Rather than a complete loss of function, the K200T mutation could produce an altered protein that exerts a dominant-negative effect. In heterozygotes, the mutant protein may compete with the wild-type subunit to form nonfunctional heterodimers, thereby disrupting the normal activity of the native BraALA3a protein complex, a common mechanism for multimetric membrane proteins whose activity depends on proper oligomerization.^53,54^ It is also plausible that a gain-of-function mutation in *BraALA3a* could confer a novel, inhibitory activity on the protein, although this is considered less likely than a dominant-negative mechanism.^55^ It is also possible that the phenotypic outcome reflects allele-specific effects. While a complete knockout might be lethal or result in more severe developmental defects, the K200T missense mutation produces a specific, viable phenotype characterized by dwarfism and compact architecture. This provides a valuable novel allele for targeted improvement of plant architecture in crop breeding. The successful identification of active target sites using the protoplast-based system confirms the efficacy of the CRISPR system in this species. However, the recovery of stably transformed regenerants appears subject to the biological limitations described above. Therefore, future efforts should focus on two strategies: gene-editing the cis-acting elements within the promoters of *BraALA3a/b/c* to regulate their transcriptional levels, and employing precise base editing to modify specific residues critical for PCZ binding, thereby creating a selective loss-of-function phenotype while avoiding disruption to conserved functional sites.

In summary, this study demonstrates that exogenous application of PCZ modulates growth and development in flowering Chinese cabbage by targeting and inhibiting BraALA3a/b/c proteins, thereby revealing a previously uncharacterized mechanism of PCZ action through phospholipid flippase inhibition (Figure 8). Heterologous expression of *BraALA3a/b/c* conferred PCZ sensitivity in yeast, establishing the concept of a specific membrane protein-based response to PCZ. *In vitro* binding assays and molecular simulations confirmed high-affinity interaction between PCZ and BraALA3a/b/c, identifying key residues within the transmembrane binding pocket and establishing a novel “membrane-target” mode of action for this pesticide, PCZ likely impeded the initiation of protein conformational self-regulation mechanisms and inhibited ATPase activity by competing for binding pocket between PI4P-a lipid flippase regulator and the transmembrane domain. Genetic perturbation in planta confirmed *BraALA3a/b/c* function in PCZ response, knockdown of *BraALA3a/b/c* resulted in dwarfed plants with reduced sensitivity to PCZ, whereas its overexpression promoted plant growth and enhanced PCZ sensitivity (Figure 8). These findings offer fundamental insights into the dwarfing mechanism induced by PCZ, enable the creation of flowering Chinese cabbage varieties, and advance sustainable strategies for managing plant architecture.

## ASSOCIATED CONTENT

### Data Availability Statement

All data supporting the findings of this study are available within the paper and within its Supporting Information published online.

### Supporting Information

Statistical analysis of growth inhibition indicators in flowering Chinese cabbage treated with PCZ under different conditions; cytological effects of PCZ on tissue microstructure; phylogenetic analysis of *ALA3* genes in *Brassicaceae* species; sensitivity of *BraALA3a/b/c^atala^*^3^ to PCZ; comparative analysis of PCZ or PI4P binding sites across BraALA3a/b/c homologs; molecular dynamics (MD) simulation analysis; protein expression of P4-ATPase and SPR negative control; transient transcriptional regulation of *BraALA3a/b/c* in response to PCZ treatment; generation, and molecular characterization of *BraALA3a/b/c* overexpression lines; other phenotypic indicators of *BraALA3a/b/c-OX*; generation and molecular characterization of *BraALA3a/b/c* knockdown lines; other phenotypic indicators of *BraALA3a/b/c^KD^*; mutation types of *BraALA3a* gene in protoplasts; types of *BraALA3a* mutations through *Agrobacterium tumefaciens*-mediated genetic transformation; detailed information on ALA3s in cruciferae plants used for phylogenetic tree; primers used in this study; and SPR binding affinities between PCZ and BraALA3a/b/c homologous proteins (PDF)

## Author Contributions

F.L., H.X., Q.G., and R.C. designed the research; Q.G., Y.S., Y.Y., S.W., X.R, Z.L., D.G., Y.C., and X.W. performed experiments; Q.G., Y.S., and Y.Y. analyzed the data; Q.G. and R.C. wrote the manuscript; F.L. and H.X. revised the manuscript. All authors have read the manuscript.

## Notes

The authors declare no competing financial interest.

## ACKNOWLEDGMENTS

We sincerely thank Takashi Akihiro (Faculty of Life and Environmental Science, Shimane University) for providing the yeast stain, Guangguang Li (Guangzhou Institute of Agriculture Science) for assisting in the seed propagation of flowering Chinese cabbage., and Changming Chen (College of Horticulture, South China Agricultural University) for verifying the *BraALA3* gene sequences of flowering Chinese cabbage.

