## Supporting Information for "The BraALA3 Homologs Mediate Propiconazole-Modulated Plant Growth in flowering Chinese cabbage"

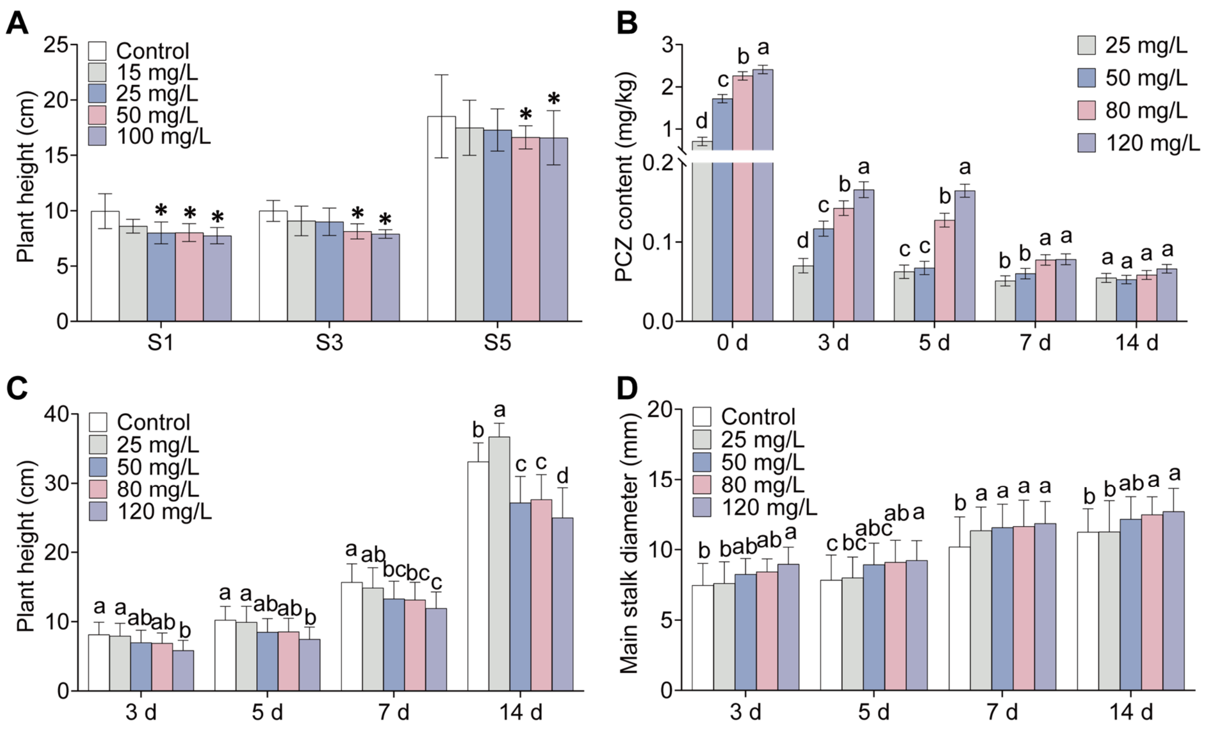


**Figure S1.** Statistical analysis of growth inhibition indicators in flowering Chinese cabbage treated with PCZ under different conditions. (A) Comparison of plant height of flowering Chinese cabbage at growth stages S1, S3, and S5 after treatment with 0, 15, 25, 50, and 100 mg/L PCZ for 7 days. Based on bolting, stalk development can be classified into five stages: S1 (seedling stage), S2 (leaf-growing stage), S3 (bolting stage), S4 (budding stage), and S5 (harvesting stage), among which S1, S3, and S5 are representative growth stages [23]. Data are mean ± SD ($n$ = 10). Significance differences were tested by two-way ANOVA (Dunnett, *$P$ < 0.05, no mark presents no significance). (B) Comparison of 0, 3, 5, 7, and 14 d PCZ absorption of PCZ-treated flowering Chinese cabbage leaves. Data are mean ± SD ($n$ = 3). Significance differences were tested by two-way ANOVA (Tukey). (C, D) Comparison of 3, 5, 7, and 14 d plant height (C), main stalk diameter (D) of 0, 25, 50, 80, and 120 mg/L PCZ-treated flowering Chinese cabbage. Data are mean ± SD ($n$ = 30). Significance differences were tested by two-way ANOVA (Tukey). Different letters in the same group data represent significant differences ($P$ < 0.05).

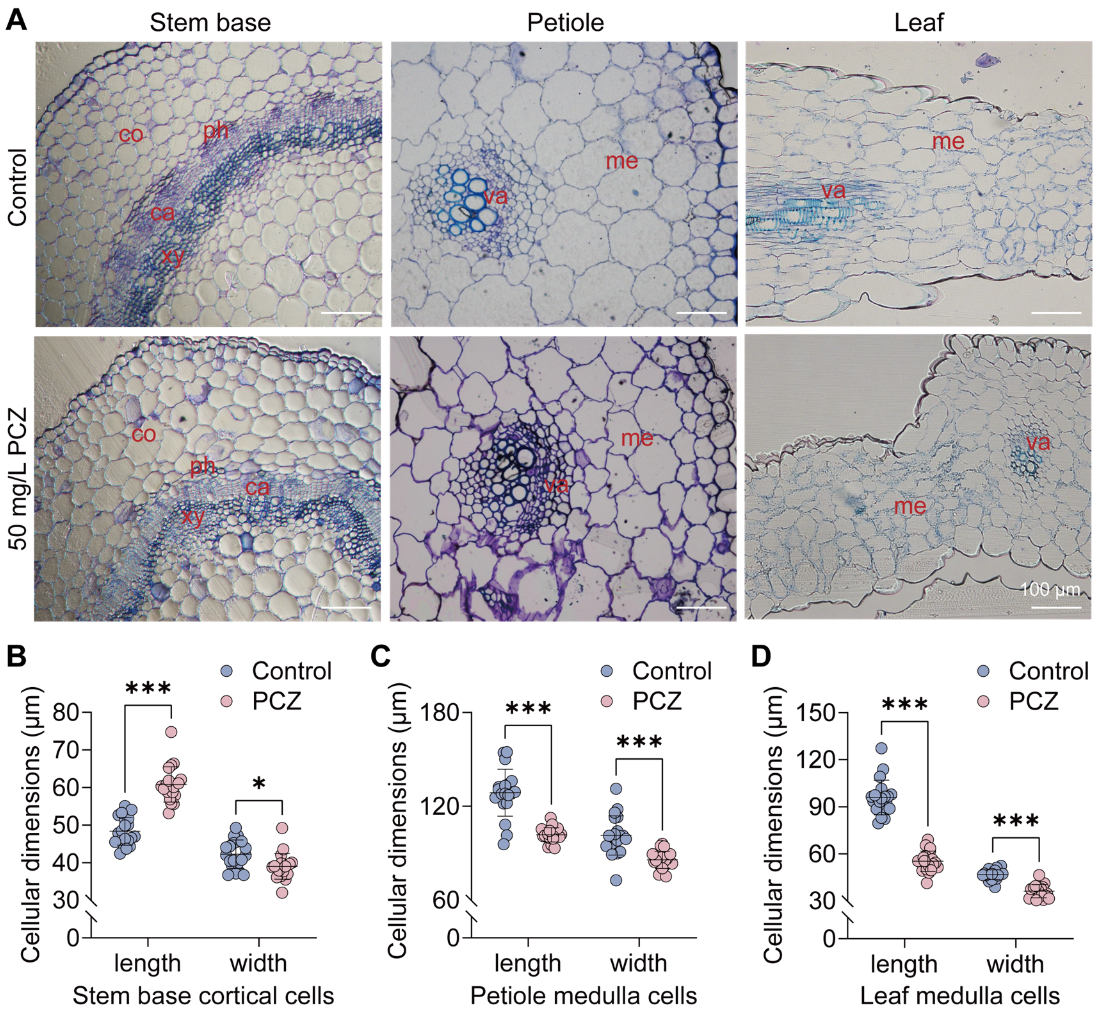


**Figure S2.** Cytological effects of PCZ on tissue microstructure in flowering Chinese cabbage. (A) Tissue sections of stem base, petiole, and leaf with control-treated or 50 mg/L PCZ-treated flowering Chinese cabbage. Bars = 100 μm. xy-xylem, ca-cambium, ph-phloem, co-cortex, va-vascular bundle, me-medulla. (B) Comparison of cortical cells length and width of stem base with control-treated or 50 mg/L PCZ-treated flowering Chinese cabbage. (C) Comparison of medulla cells length and width of petiole with control-treated or 50 mg/L PCZ-treated flowering Chinese cabbage. (D) Comparison of medulla cells length and width of leaf with control-treated or 50 mg/L PCZ-treated flowering Chinese cabbage. Data are mean ± SD ($n$ = 20). Significance differences were tested by two-way ANOVA (Šídák, $*P < 0.05$, $**P < 0.01$, $***P < 0.001$).

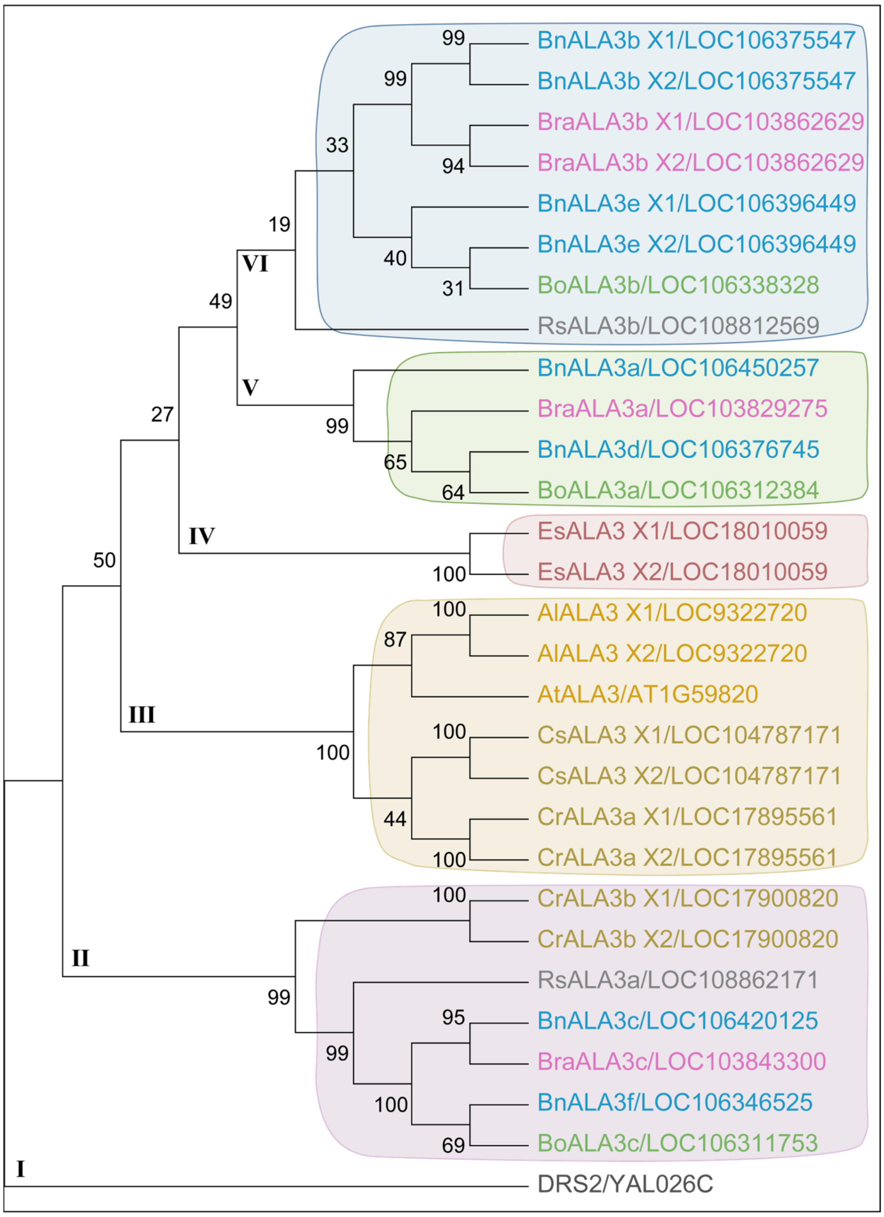


**Figure S3.** Phylogenetic analysis of *ALA3* genes in *Brassicaceae* species. Phylogenetic relationships analyzed by neighbor-joining method. Multiple sequence alignment was performed using ClustalW in MEGA 11.0 with default parameters. Bootstrap values from 1,000 replicates are shown at branch nodes. Scale bar indicates 0.1 substitutions per nucleotide position. Sc-*Saccharomyces cerevisiae*, Al-*Arabidopsis lyrata subsp. lyrate*, At-*Arabidopsis thaliana*, Bn-*Brassica napus*, Bo-*Brassica oleracea var. oleracea*, Bra-*Brassica rapa*, Cs-*Camelina sativa*, Cr-*Capsella rubella*, Es-*Eutrema salsugineum*, Rs-*Raphanus sativus*.

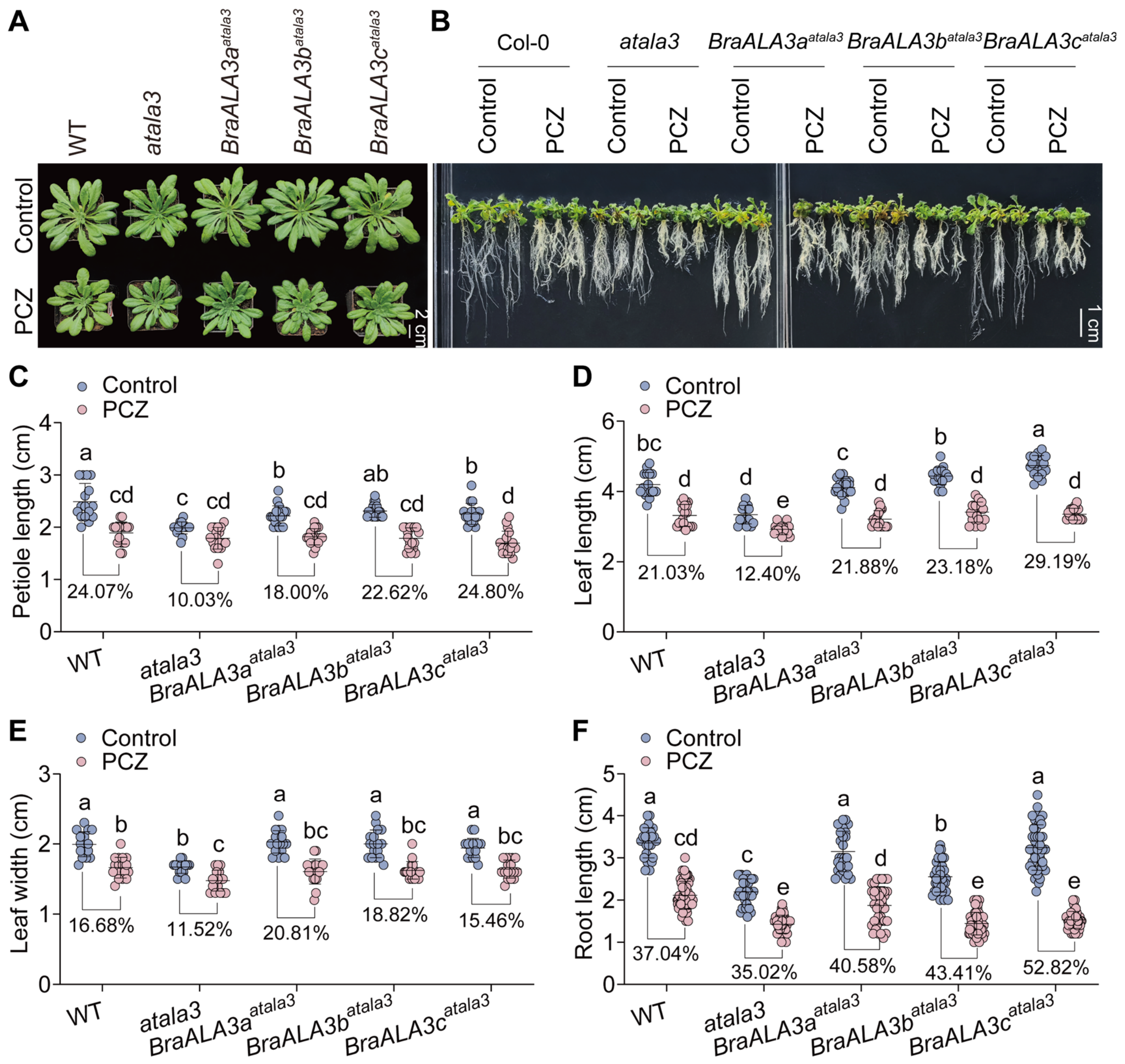


**Figure S4.** Sensitivity of *BraALA3a/b/c*$^{atala3}$ to PCZ. (A) 5 d phenotypes of control-treated or 50 mg/L PCZ-treated WT, *atala3*, and *BraALA3a/b/c*$^{atala3}$ (28-day-old). Bar = 2 cm. (B) 5 d phenotypes of WT, *atala3*, and *BraALA3a/b/c*$^{atala3}$ grown on 1/2 MS medium containing DMSO (control) or 1 μM PCZ. Bar = 1 cm. (C-E) Comparison of 5 d petiole length (C), leaf length (D), and leaf width (E) of control-treated or 50 mg/L PCZ-treated WT, *atala3*, and *BraALA3a/b/c*$^{atala3}$. Data are mean ± SD ($n \geq 16$). (F) Comparison of 5 d root length of WT, *atala3*, and *BraALA3a/b/c*$^{atala3}$ grown on 1/2MS medium containing DMSO (control) or 1 μM PCZ. Data are mean ± SD ($n \geq 28$). Significance differences were tested by one-way ANOVA (Tukey). Different letters in the same group data represent significant differences ($P < 0.05$). Percentages represent the decrease or increase of each indicator.

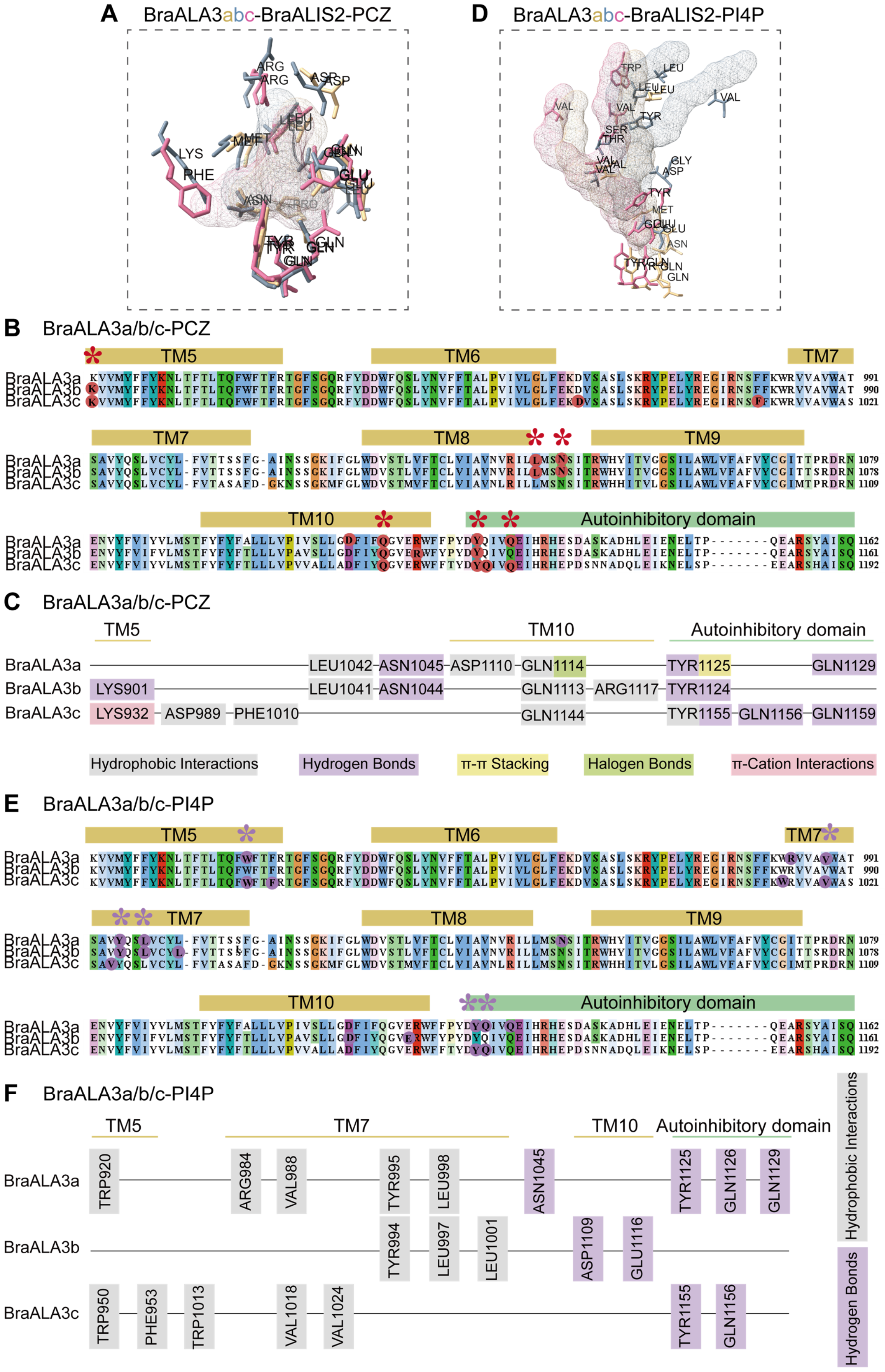


**Figure S5.** Comparative analysis of PCZ or PI4P binding sites across BraALA3a/b/c homologs. (A) Molecular docking of homologous proteins BraALA3a/b/c with the same ligand PCZ. Ligands are represented by the mesh model. Golden, blue, and pink stick model represent BraALA3a, BraALA3b, and BraALA3c amino acid residue, respectively. (B) The presentation of primary sequences at molecular docking sites between homologous proteins BraALA3a/b/c and PCZ. Colored sequences represent regions with high conservatism. The box above the sequence is marked as three-level structure. Red star labeled as docking site with PCZ. (C) Schematic of primary sequences at molecular docking sites between homologous proteins BraALA3a/b/c and PCZ. The binding of BraALA3a/b/c homologous proteins to PCZ is stabilized by hydrophobic interactions (gray), hydrogen bonds (purple), $\pi$-$\pi$ stacking (yellow), halogen bonds (green), and $\pi$-cation interactions (pink). (D) Molecular docking of homologous proteins BraALA3a/b/c with the same ligand PI4P. (E) The presentation of primary sequences at molecular docking sites between homologous proteins BraALA3a/b/c and PI4P. Purple star labeled as docking site with PI4P. (F) Schematic of primary sequences at molecular docking sites between homologous proteins BraALA3a/b/c and PI4P. The binding of BraALA3a/b/c homologous proteins to PI4P is stabilized by hydrophobic interactions (gray) and hydrogen bonds (purple).

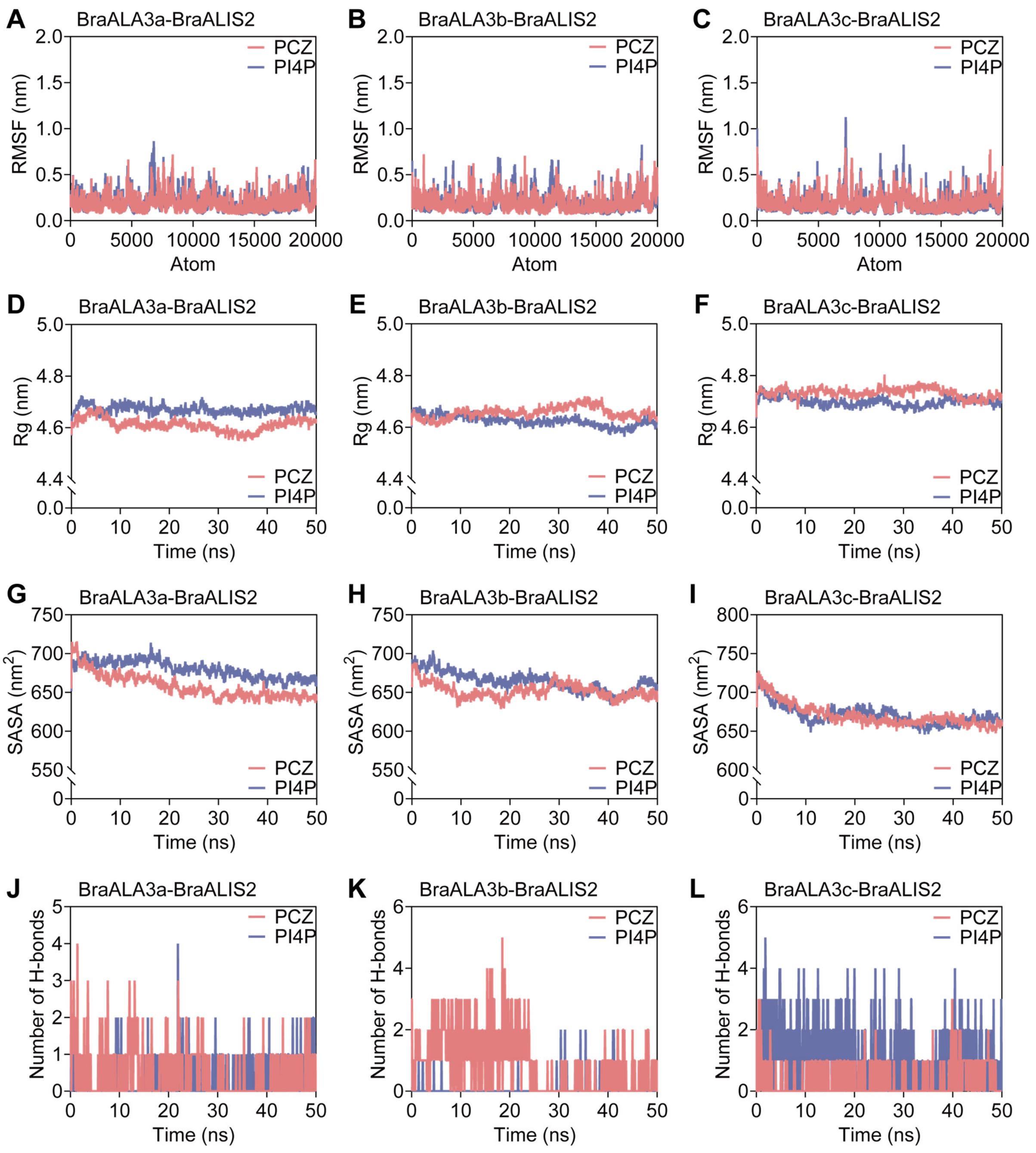


**Figure S6.** Molecular dynamics (MD) simulation analysis of BraALA3a/b/c-BraALIS2 protein binding to PCZ/PI4P. (A-C) The Root Mean Square Fluctuation (RMSF) curves of protein skeleton atoms in BraALA3a/b/c-BraALIS2 and PCZ/PI4P complexes. (D-F) The Radius gyration (Rg) curves of BraALA3a/b/c-BraALIS2 and PCZ/PI4P complexes over time. (G-I) The Solvent Accessible Surface Area (SASA) curves of BraALA3a/b/c-BraALIS2 and PCZ/PI4P complexes over time. (J-L) The hydrogen bonds number curves of BraALA3a/b/c-BraALIS2 and PCZ/PI4P complexes over time.

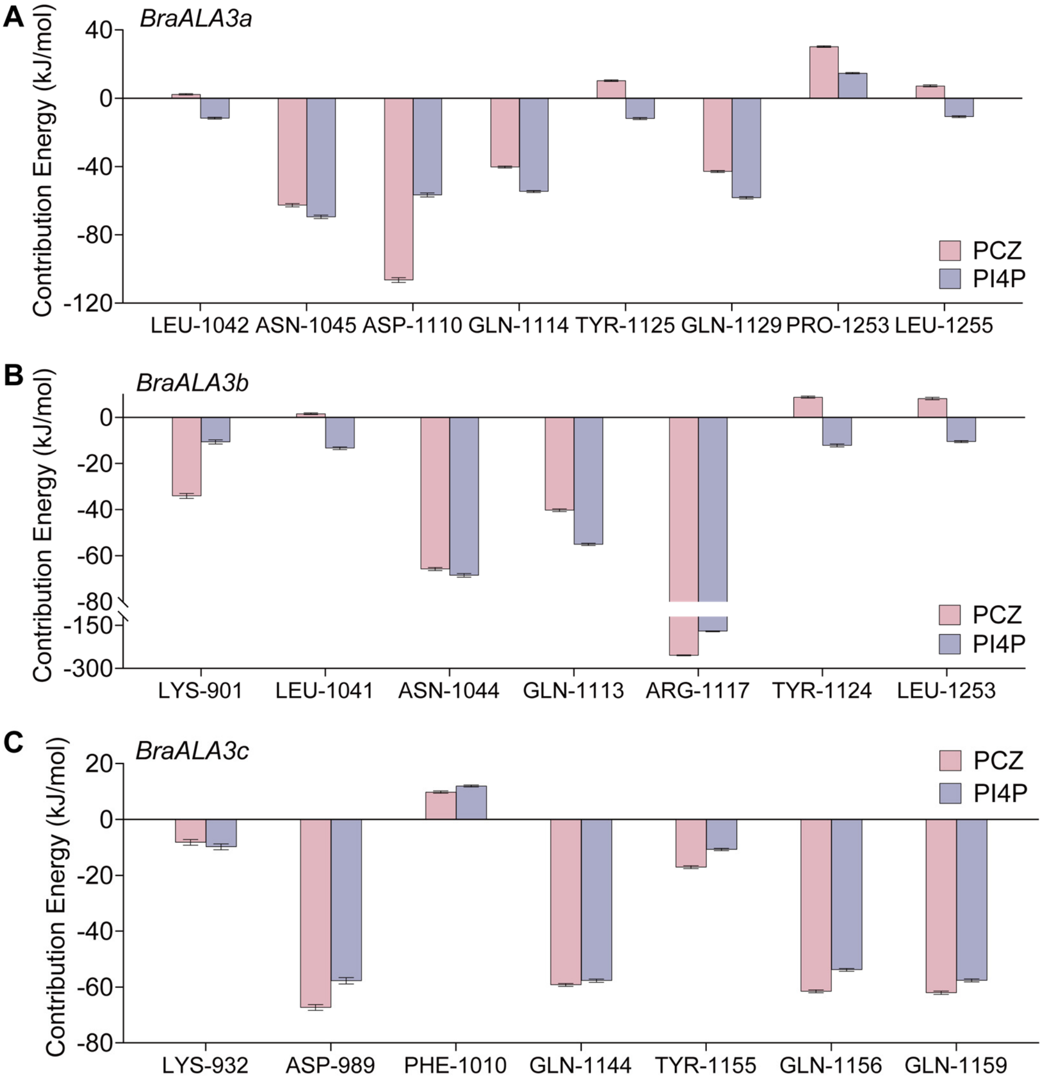


**Figure S7.** Free energy contribution of key amino acids residues in BraALA3a/b/c-BraALIS2-PCZ/PI4P protein-substrate complex.

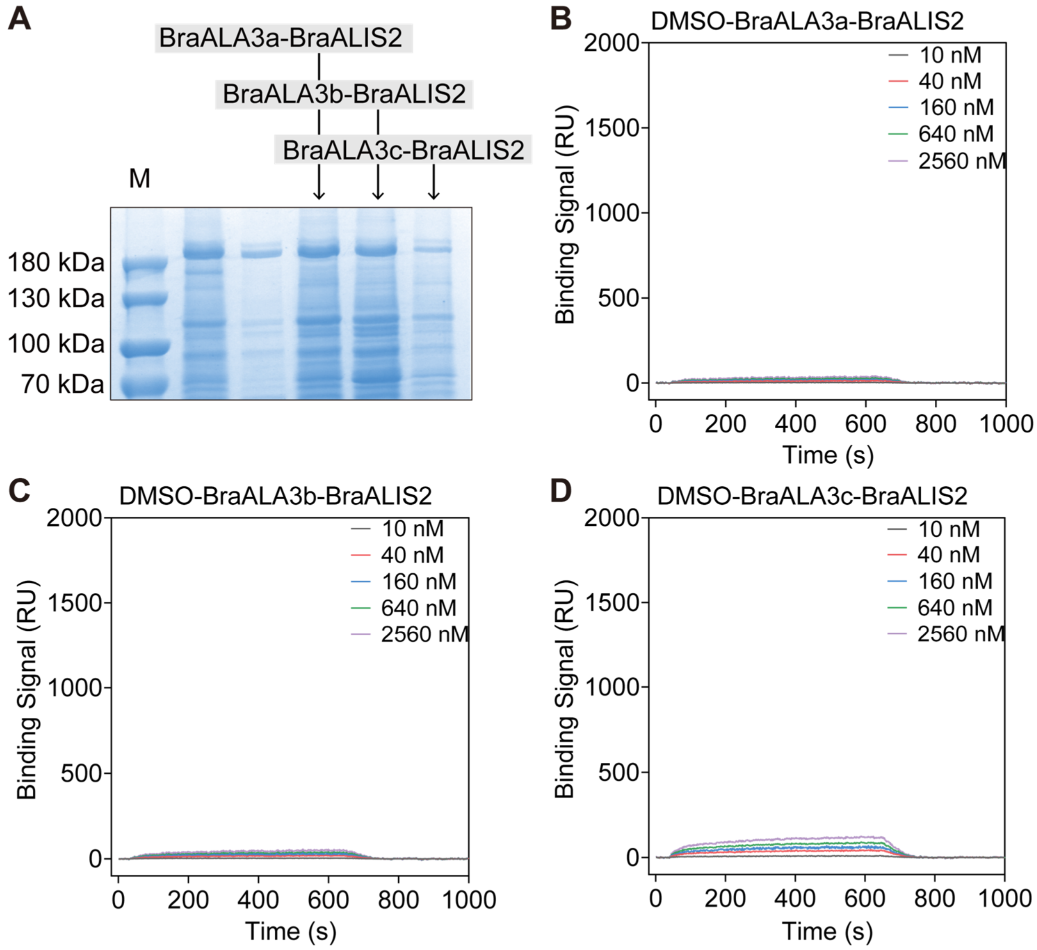


**Figure S8.** Protein expression of P4-ATPase and SPR negative control. (A) SDS-PAGE of P4-ATPase protein expression. (B-D) SPR affinity binding signals of negative control.

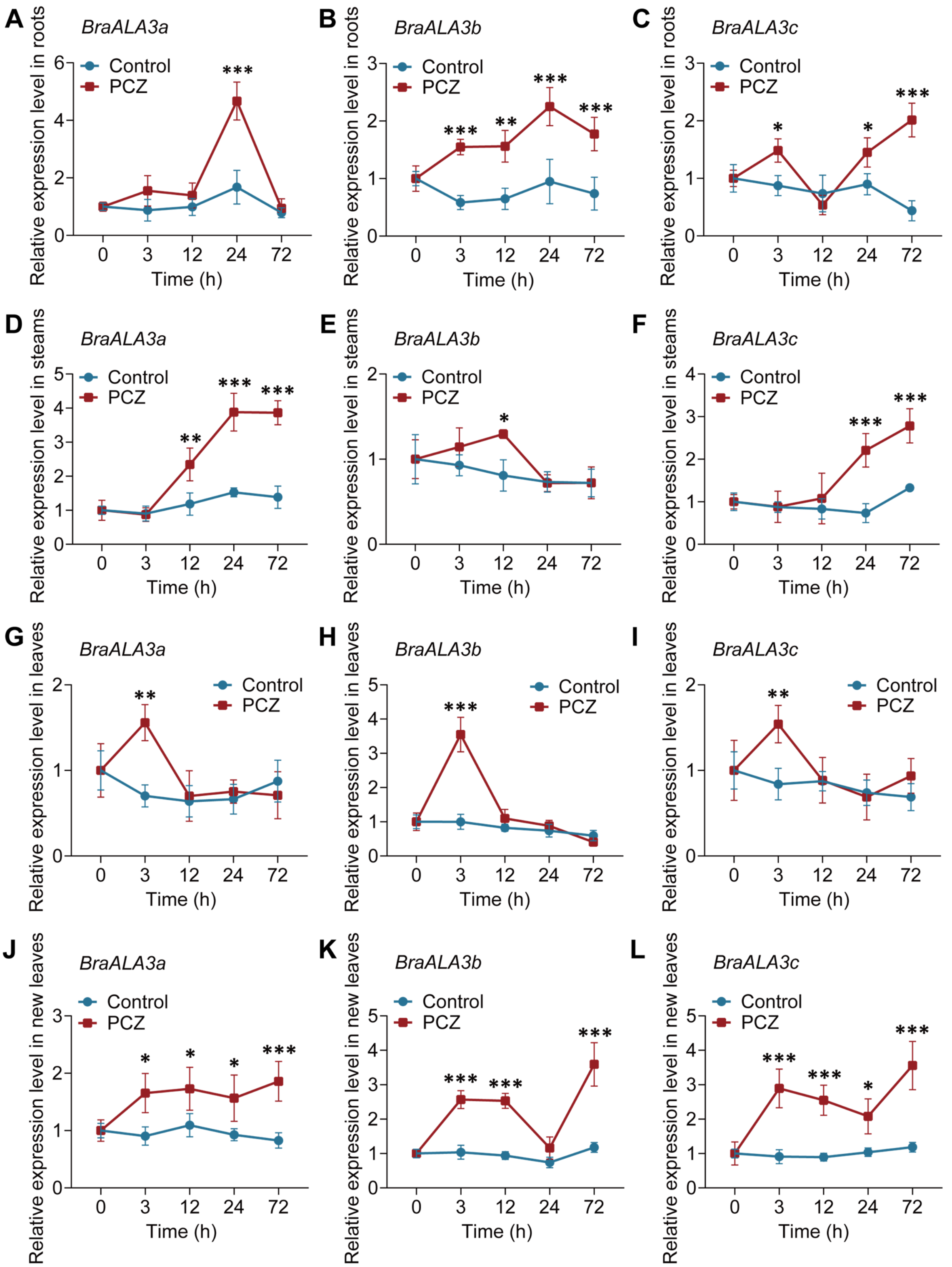


**Figure S9.** Transient transcriptional regulation of *BraALA3a/b/c* in response to PCZ treatment. Temporal dynamics of *BraALA3a/b/c* expression in PCZ-treated flowering Chinese cabbage roots (A-C), steams (D-F), leaves (G-I), and new leaves (J-L). Relative transcript levels measured by qRT-PCR (mean ± SD, $n$ = 3). Temporal expression profiles normalized to untreated controls (0 h). Significance differences were tested by Two-way ANOVA (Šídák, *$P < 0.05$, **$P < 0.01$, ***$P < 0.001$, no mark presents no significance).

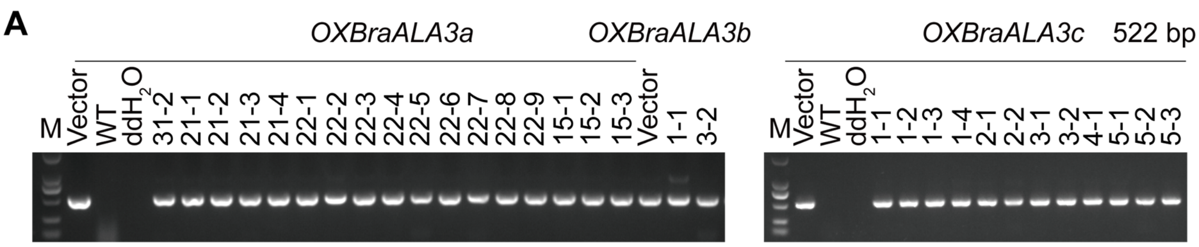


**B**

| | *BraALA3a* | *BraALA3b* | *BraALA3c* |
|---|---|---|---|
| WT | 1.02 | 1.01 | 1.01 |
| *OXBraALA3a31-2* | 9.75 | 7.73 | 5.25 |
| *OXBraALA3a21-1* | 6.27 | 5.16 | 2.78 |
| *OXBraALA3a21-2* | 3.32 | 3.92 | 3.60 |
| *OXBraALA3a21-3* | 14.30 | 4.51 | 5.59 |
| *OXBraALA3a21-4* | 3.06 | 4.54 | 3.74 |
| *OXBraALA3a22-1* | 7.70 | 2.34 | 3.24 |
| *OXBraALA3a22-2* | 4.84 | 1.63 | 17.56 |
| *OXBraALA3a22-3* | 6.63 | 3.24 | 26.93 |
| *OXBraALA3a22-4* | 3.96 | 1.76 | 36.77 |
| *OXBraALA3a22-5* | 2.56 | 1.71 | 17.91 |
| *OXBraALA3a22-6* | 4.71 | 3.15 | 10.76 |
| *OXBraALA3a22-7* | 2.56 | 1.75 | 14.31 |
| *OXBraALA3a22-8* | 2.74 | 2.44 | 3.97 |
| *OXBraALA3a22-9* | 4.57 | 3.34 | 10.07 |
| *OXBraALA3a15-1* | 3.92 | 2.77 | 4.62 |
| *OXBraALA3a15-2* | 4.13 | 2.13 | 7.31 |
| *OXBraALA3a15-3* | 7.64 | 2.30 | 2.84 |

30 20 10

**C**

| | *BraALA3b* | *BraALA3a* | *BraALA3c* |
|---|---|---|---|
| WT | 1.01 | 1.01 | 1.00 |
| *OXBraALA3b1-1* | 12.81 | 58.17 | 30.03 |
| *OXBraALA3b3-2* | 17.12 | 19.50 | 58.56 |

40 20

**D**

| | *BraALA3c* | *BraALA3a* | *BraALA3b* |
|---|---|---|---|
| WT | 1.02 | 1.00 | 1.00 |
| *OXBraALA3c1-1* | 6.03 | 6.23 | 37.56 |
| *OXBraALA3c1-2* | 5.59 | 4.43 | 29.11 |
| *OXBraALA3c1-3* | 2.00 | 7.63 | 19.53 |
| *OXBraALA3c1-4* | 11.59 | 19.76 | 56.38 |
| *OXBraALA3c2-1* | 2.99 | 9.88 | 31.22 |
| *OXBraALA3c2-2* | 4.38 | 12.65 | 86.91 |
| *OXBraALA3c3-1* | 6.63 | 7.15 | 33.47 |
| *OXBraALA3c3-2* | 5.76 | 9.03 | 66.13 |
| *OXBraALA3c4-1* | 1.40 | 7.53 | 30.39 |
| *OXBraALA3c5-1* | 3.40 | 5.39 | 3.42 |
| *OXBraALA3c5-2* | 2.64 | 14.78 | 4.13 |
| *OXBraALA3c5-3* | 3.41 | 5.59 | 2.41 |

80604020

**Figure S10.** Generation, and molecular characterization of *BraALA3a/b/c* overexpression lines in flowering Chinese cabbage. (A) Identification of *BraALA3a/b/c* overexpression lines by PCR. (B-D) Identification of *BraALA3a/b/c* overexpression lines by qRT-PCR. Data are mean ± SD ($n$ = 3). Temporal expression profiles normalized to WT.

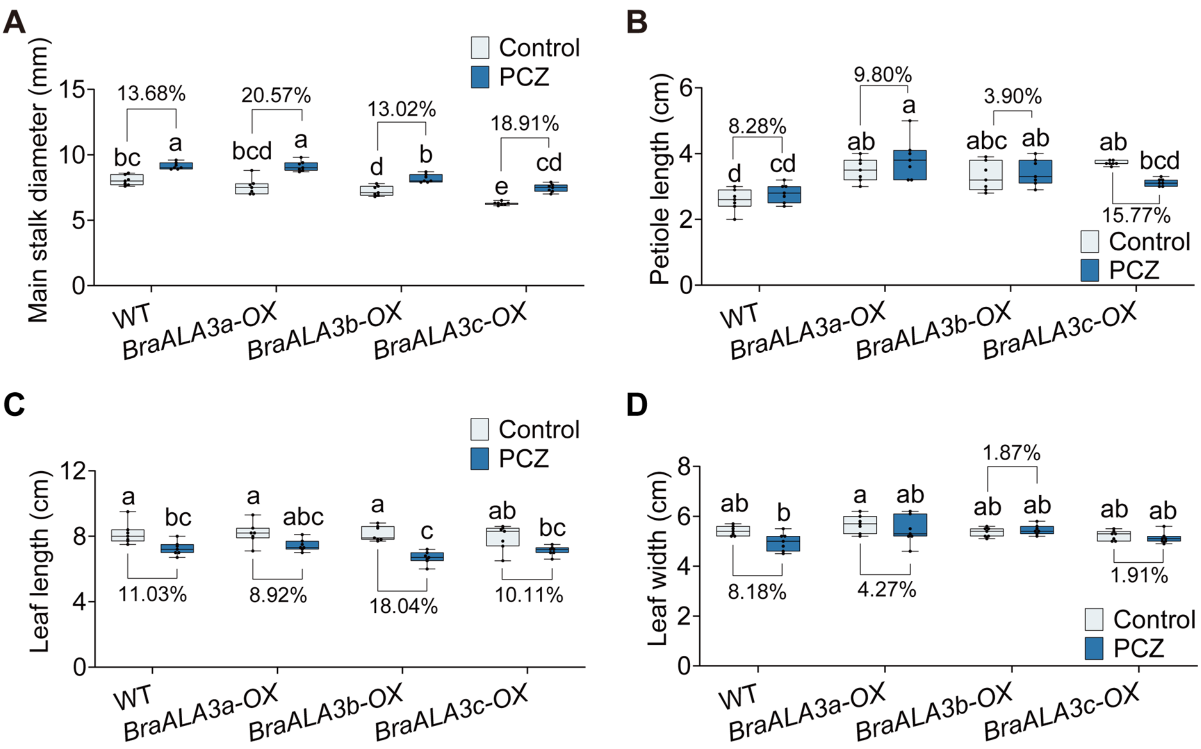


**Figure S11.** Other phenotypic indicators of *BraALA3a/b/c-OX*. Comparison of 5 d main stalk diameter (A), deformed leaf petiole length (B), Leaf length (C), and Leaf width (D) of control-treated or 50 mg/L PCZ-treated WT and *BraALA3a/b/c-OX*. Data are mean ± SD ($n$ = 7). Significance differences were tested by one-way ANOVA (Tukey). Different letters represent significant differences ($P < 0.05$). Percentages represent the decrease or increase of each indicator.

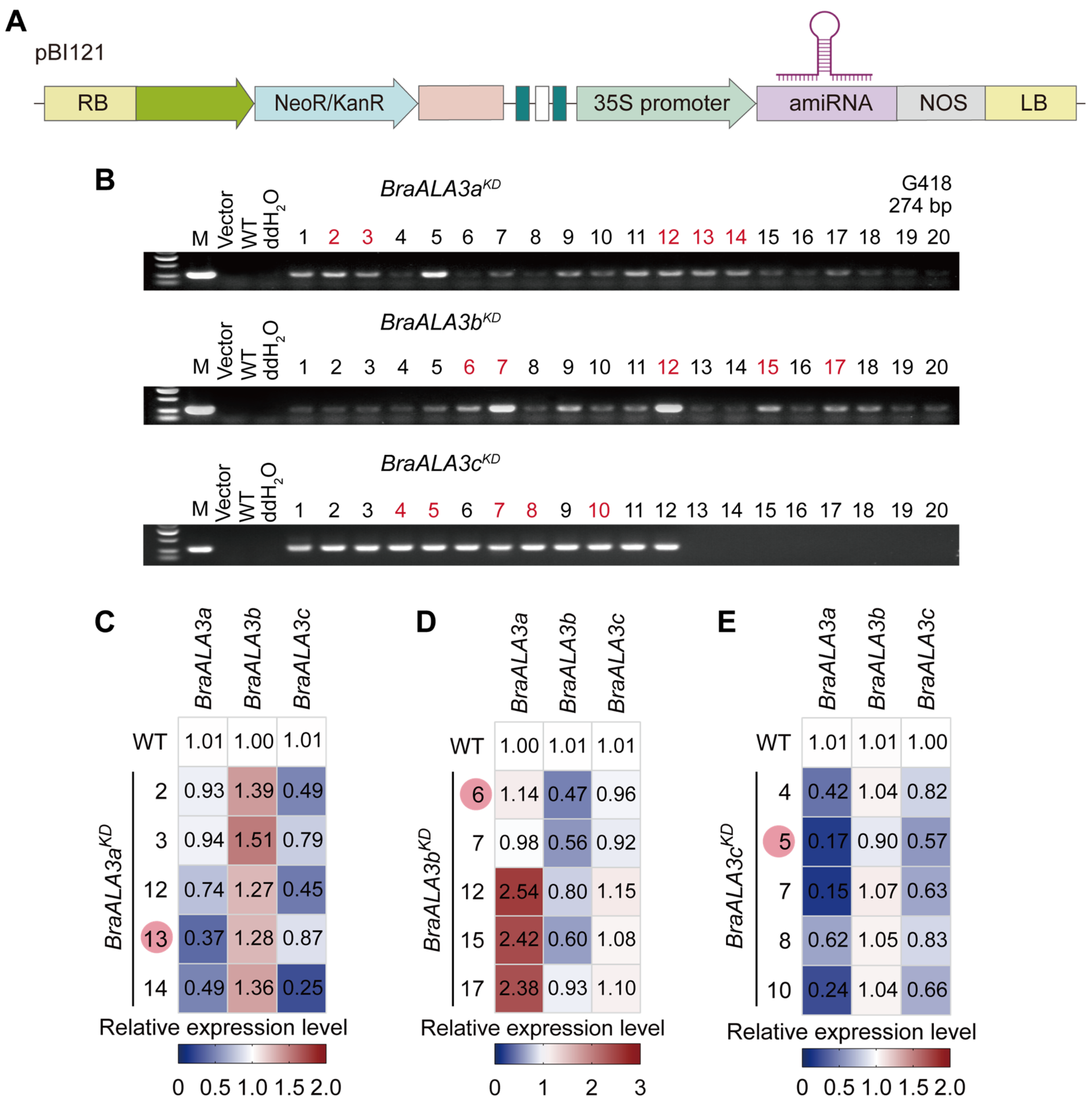


**Figure S12.** Generation and molecular characterization of *BraALA3a/b/c* knockdown lines in flowering Chinese cabbage. (A) Vector of floral dip transformation for generating *BraALA3a/b/c* knockdown lines in flowering Chinese cabbage. (B) Identification of *BraALA3a/b/c* knockdown lines by PCR. Plants labeled in red were selected for further analysis of gene expression. (C-E) Identification of *BraALA3a/b/c* knockdown lines by qRT-PCR. Data are mean ± SD ($n$ = 3). Temporal expression profiles normalized to WT. Plants indicated by red circles were selected for phenotypic analysis.

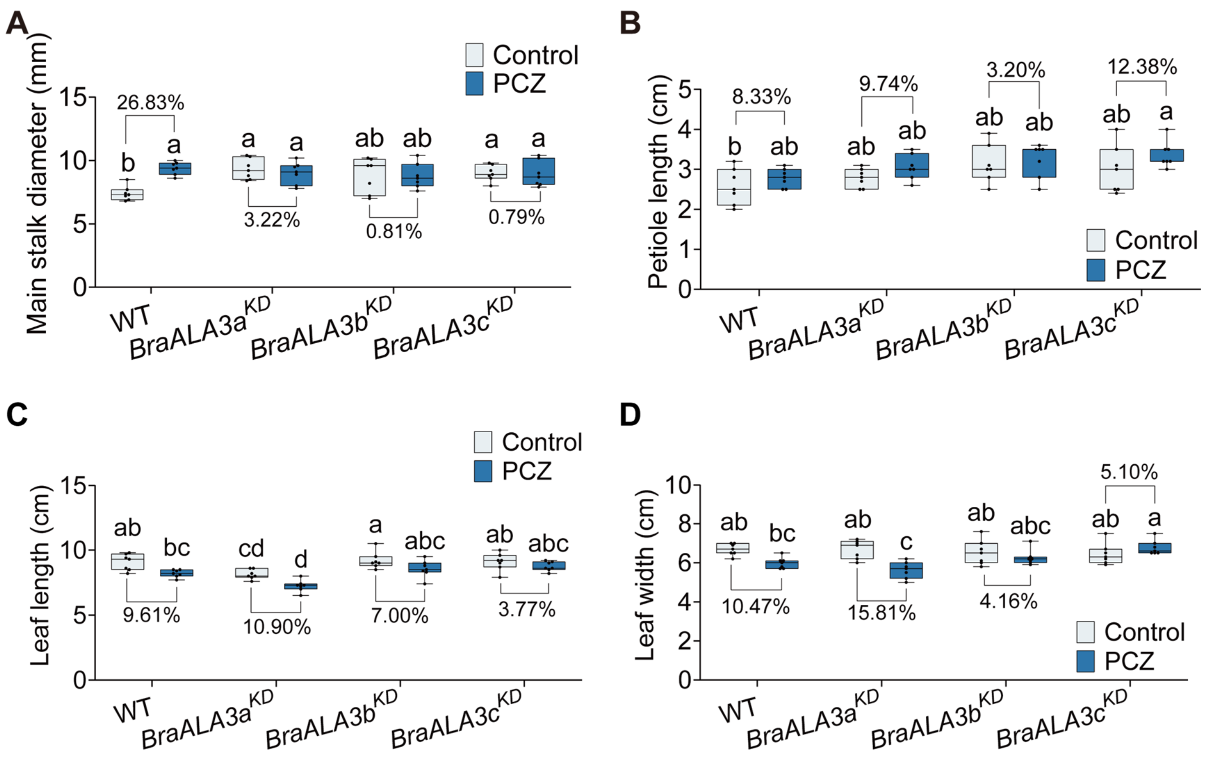


**Figure S13.** Other phenotypic indicators of *BraALA3a/b/c*$^{KD}$. Comparison of 5 d main stalk diameter (A), deformed leaf petiole length (B), Leaf length (C), and Leaf width (D) of control-treated or 50 mg/L PCZ-treated WT and *BraALA3a/b/c*$^{KD}$. Data are mean ± SD ($n$ = 7). Significance differences were tested by one-way ANOVA (Tukey). Different letters represent significant differences ($P$ < 0.05). Percentages represent the decrease or increase of each indicator.

**A** *BraA06.GA4* — Encoding area

PAM / sgRNA

| Sequence | |
|---|---|
| ATGTGGTCGGAAGGTTTCACCATCACCGGCTCCCCTCTCAACGACTTCCGTAAACTTTGG | Wild Type |
| ATGTGGTCGGAAGGTTTCACCATCA- -GGCTCCCCTCTCAACGACTTCCGTAAACTTTGG | **434 (-2 bp)** |
| ATGTGGTCGGAAGGTTTCACCATCA- - -GCTCCCCTCTCAACGACTTCCGTAAACTTTGG | **434 (-3 bp)** |
| ATGTGGTCGGAAGGTTTCACCATCA**A**CCGGCTCCCCTCTCAACGACTTCCGTAAACTTTGG | **434 (+1 bp)** |
| ATGTGGTCGGAAGGTTTCACCATAACCGGCTCCCCTCTCAACGACTTCCGTAAACTTTGG | 432C>A |

| Protein | |
|---|---|
| MWSEGFTITGSPLNDFRKLW | Wild Type |
| MWSEGFTI**RLPSQRLP*** | **145 (frameshift mutation) / 153 (Early termination of base translation)** |
| MWSEGFTI**S**-SPLNDFRKLW | **145T>S /146 -1aa** |
| MWSEGFTI**NRLPSQRLP*** | **145 (frameshift mutation) / 154 (Early termination of base translation)** |

**B** *BraALA3a*-sgRNA02

PAM / sgRNA

| Sequence | |
|---|---|
| ATCCAGCCTCAGGCTCCGTCTTATCGAACCGTCTACTGCAATGATCGAGATTCCAATATG | Wild Type |
| ATACAGCCTCAGGCTCCGTCTTATCGAACCGTCTACTGCAATGATCGAGATTCCAATATG | 87C>A |
| ATCCAGCCTCAGGCTCCGTCTT**G**TCGTACCGTCTACTGCAATGATCGAGATTCCAATATG | **107A>G** 111A>T |
| ATACAGCCTCAGGCTCCGTCTT**G**TCGTACCGTCTACTGCAATGATCGAGATTCCAATATG | 87C>A **107A>G** 111A>T |
| ATCCAGCCTCAGGCTCCGTCTTATCGAACCGTCTACTGCAATGATCGAGATTCCAACATG | 141T>C |

| Protein | |
|---|---|
| IQPQAPSYRTVYCNDRDSNM | Wild Type |
| IQPQAPS**C**RTVYCNDRDSNM | **36Y>C** |

**C** *BraALA3a*-sgRNA07

sgRNA / PAM

| Sequence | |
|---|---|
| TGTGGTGTCACTGAAATAGAAAGAGGAATCGCTCAGCGTAATGGCTTAAAAGTACACGAG | Wild Type |
| TGTGGTGTCACTGAAATAGAAA**A**AGGAATCGCTCAGCGTAATGGCTTAAAAGTACACGAG | **1334G>A** |
| TGTGGTGTCACTGAAATAGAAAGGGGAATCGCTCAGCGTAATGGCTTAAAAGTACACGAG | 1335A>G |
| TGTGGTGTCACTGAAATAGAAAGAGGGATCGCTCAGCGTAATGGCTTAAAAGTACACGAG | 1338A>G |

| Protein | |
|---|---|
| CGVTEIERGIAQRNGLKVHE | Wild Type |
| CGVTEIE**K**GIAQRNGLKVHE | 445R>K |

**Figure S14.** Mutation types of *BraALA3a* gene in protoplasts through comparative sequencing analysis. (A) Positive control: types of *BraA06.GA4* gene mutations in flowering Chinese cabbage protoplasts. (B, C) Types of *BraALA3a*-sgRNA02 (B) and *BraALA3a*-sgRNA07 (C) mutations in flowering Chinese cabbage protoplasts. PAM sites are highlighted in pink, sgRNA are highlighted in blue, mutated nucleotides were highlighted in red.

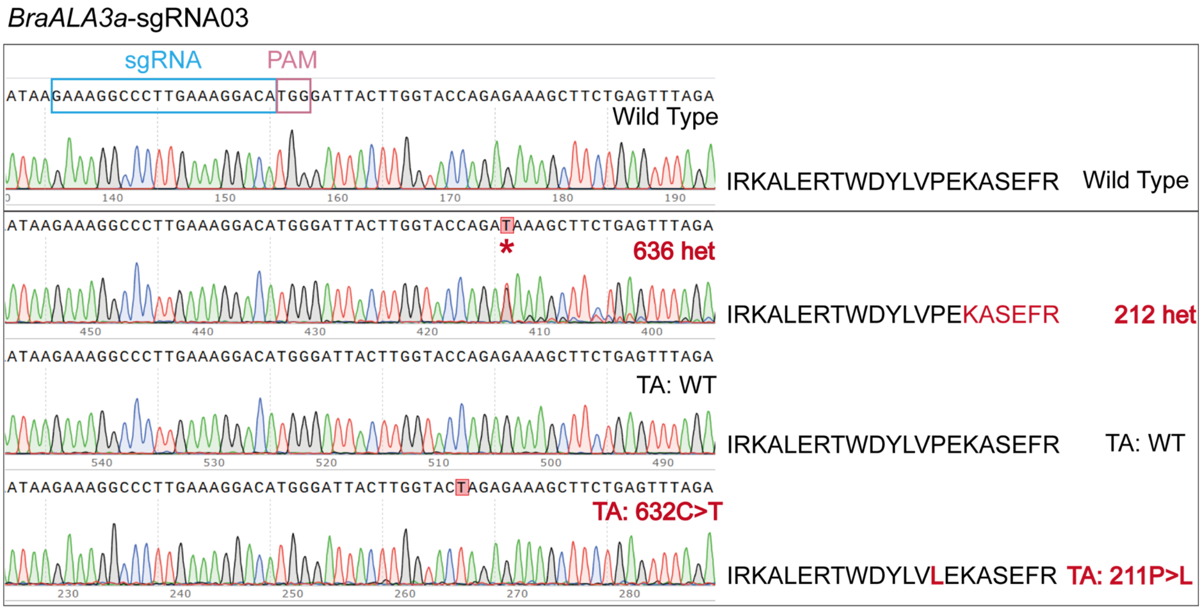


**Figure S15.** Types of *BraALA3a* mutations in flowering Chinese cabbage through *Agrobacterium tumefaciens*-mediated genetic transformation. PAM sites are highlighted in pink, sgRNA are highlighted in blue, mutated nucleotides were highlighted in red.

**Table S1.** Detailed information on ALA3s in cruciferae plants used for phylogenetic tree.

| Species | Gene name | Gene ID |
|---|---|---|
| *Arabidopsis lyrata* | AlALA3 X1/X2 | LOC9322720 |
| *Arabidopsis thaliana* | AtALA3 | AT1G59820 |
| *Brassica napus* | BnALA3a X1/X2 | LOC106396449 |
| | BnALA3b X1/X2 | LOC106375547 |
| | BnALA3c X1/X2 | LOC106420125 |
| | BnALA3d | LOC106376745 |
| | BnALA3e | LOC106450257 |
| | BnALA3f | LOC106346525 |
| *Brassica oleracea* | BoALA3a | LOC106338328 |
| | BoALA3b | LOC106342273 |
| | BoALA3c | LOC106342274 |
| | BoALA3d | LOC106312384 |
| | BoALA3e | LOC106311753 |
| *Brassica rapa* | BraALA3a | LOC103829275 |
| | BraALA3b X1/X2 | LOC103862629 |
| | BraALA3c | LOC103843300 |
| *Camelina sativa* | CsALA3 X1/X2 | LOC104787171 |
| *Capsella rubella* | CrALA3a X1/X2 | LOC17895561 |
| | CrALA3b X1/X2 | LOC17900820 |
| *Eutrema salsugineum* | EsALA3 X1/X2 | LOC18010059 |
| *Raphanus sativus* | RsALA3a | LOC108812569 |
| | RsALA3b | LOC108862171 |
| | RsALA3c | LOC108807803 |
| *Saccharomyces cerevisiae* | DRS2 | YAL026C |

**Table S2.** Primers used in this study.

| Primer name | Primer sequence (5' to 3') |
|---|---|
| **Yeast expression** | |
| pYES2-BraALA3a-F | gctgtaatacgactcactatagggaatattaagcttATGGTTCGATCGGGTAATTTAAGCG |
| pYES2-BraALA3a-R | gcatgctcgagcggccgccagtgtgatggatatctgcagaattcCTACTTCTTCTTTGGTACTGTCGGCC |
| pYES2-BraALA3b-F | gctgtaatacgactcactatagggaatattaagcttATGGTACGATCGGGTAGTTTAAACGG |
| pYES2-BraALA3b-R | gcatgctcgagcggccgccagtgtgatggatatctgcagaattcCTACTTCTTTGGTACTTTAGGGCGTGAC |
| pYES2-BraALA3c-F | gctgtaatacgactcactatagggaatattaagcttATGGCTAGCTCCGGAGGTGGATT |
| pYES2-BraALA3c-R | gcatgctcgagcggccgccagtgtgatggatatctgcagaattcTCATTTCTTCTTTGGTGCTT |
| pYES2-357-F | TTTCGGTTTGTATTACTTCTTATTC |
| BraALA3a-R | CGGCTGGAAAAAACGCATCTTTTTAATC |
| BraALA3b-R | CTGCTGGAAAAAGGCATCTTGTTTTATC |
| BraALA3c-R | GTTGATCTTGCAGAACCTCCACTGTACTG |
| **Protein expression in vitro** | |
| 424-RA-R | TGGACCTTGAAACAAAACTTCCAA |
| 424-RA-F | CTCGAGTCATGTAATTAGTTATGTC |
| pRS424-BraALIS2-F | aagttttgtttcaaggtccaggcgcgcctATGATGGAAGTGGAAGGATC |
| pRS424-BraALIS2-R | aactaattacatgactcgagggcgcgcctTCAACTCGAAAGGCTTTTCTT |
| BraALIS2-R | TCTTCAGGAATGCACTCAAC |
| 426-RA-R | TGGACCTTGAAACAAAACTTCCAACTT |
| 426-RA-F | CTCGAGTCATGTAATTAGTTATGTCACG |
| pRS426-BraALA3a-F | gaaaagttggaagttttgtttcaaggtccaATGGTTCGATCGGGTAATTTAAGCG |
| pRS426-BraALA3a-R | cgtgacataactaattacatgactcgagCTACTTCTTCTTTGGTACTGTCGGC |
| pRS426-BraALA3b-F | gaaaagttggaagttttgtttcaaggtccaATGGTACGATCGGGTAGTTTAAACG |
| pRS426-BraALA3b-R | cgtgacataactaattacatgactcgagCTACTTCTTTGGTACTTTAGGGCGT |
| pRS426-BraALA3c-F | gaaaagttggaagttttgtttcaaggtccaATGGCTAGCTCCGGAGGTG |
| pRS426-BraALA3c-R | cgtgacataactaattacatgactcgagTCATTTCTTCTTTGGTGCTTTGGGT |

| Primer name | Primer sequence (5' to 3') |
| --- | --- |
| **RT-qPCR** | |
| qRT-BraUBC10-F | GGGTCCTACAGACAGTCCTTAC |
| qRT-BraUBC10-R | ATGGAACACCTTCGTCCTAAA |
| qRT-BraALA3a-F | ACACCACGTGATAGAAATGAAAATG |
| qRT-BraALA3a-R | CGCCTAGAAGAGAAACAATGGG |
| qRT-BraALA3b-F | CGATTTCATCTACCAAGGGGTT |
| qRT-BraALA3b-R | TGCTTTGAGAGCTCCCGT |
| qRT-BraALA3c-F | CGCCACGTGATAGAAACGAGA |
| qRT-BraALA3c-R | GAAGAACCACCTCTCCACCC |
| **Gene silencing** | |
| pre-miRNA-F | ggactctagaggatccGGGTGAGAATCTCCATGT |
| pre-miRNA-R | gatcggggaaattcgagctcGGGTGAAGAGCTCATGT |
| G418-F | CGGCTATGACTGGGCACAACAGACAAT |
| G418-R | CTCGGCAGGAGCAAGGTGAGATGAC |
| **Gene overexpression** | |
| pBI121-BraALA3a-F | gagagaacacgggggactctagaATGGTTCGATCGGGTAATTTAA |
| pBI121-BraALA3a-R | aacgatcggggaaattcgagctcCTACTTCTTCTTTGGTACTGTC |
| pBI121-BraALA3b-F | gagagaacacgggggactctagaATGGTACGATCGGGTAGTTTAA |
| pBI121-BraALA3b-R | aacgatcggggaaattcgagctcCTACTTCTTTGGTACTTTAGGG |
| pBI121-BraALA3c-F | gagagaacacgggggactctagaATGGCTAGCTCCGGAGGTGGAT |
| pBI121-BraALA3c-R | aacgatcggggaaattcgagctcTCATTTCTTCTTTGGTGCTTTG |
| Neo-F | CCTGTCCGGTGCCCTGAATGAA |
| Neo-R | CGGGTAGCCAACGCTATGTCCT |
| **Gene editing** | |
| Crispr-BraA06.GA4-sg-F | cctctaatacgactcactataGGTTGAGAGGGGAGCCGGTGAgtttaagagctatgc |
| BraA06.GA4-sgRNA-F | GCCCAGTTTAAATCTGGATT |
| BraA06.GA4-sgRNA-R | AAAACATCCCTCTCATTGAC |

| Primer name | Primer sequence (5' to 3') |
|---|---|
| TF-BraA06.GA4-sgRNA | ctcggagtgatcgcacAGACAGGCTTCTCTGGCTACG |
| TR-BraA06.GA4-sgRNA | ctgagaggctggatggAGATATATACCAGTAGTTGTTAAGG |
| Crispr-BraALA3a-sg02-F | cctctaatacgactcactataGGCAGTAGACGGTTCGATAAGAgtttaagagctatgc |
| BraALA3a-sgRNA02-F | TTCGTCGTCGGCTAGTCATC |
| BraALA3a-sgRNA02-R | CGCCTAAACTGTCAGGGAAA |
| TF-BraALA3a-sgRNA02 | ctcggagtgatcgcacTTCGTCGTCGGCTAGTCATC |
| TR-BraALA3a-sgRNA02 | ctgagaggctggatggAAAGCACAAATTCCAAATGC |
| Crispr-BraALA3a-sg07-F | cctctaatacgactcactataGGAGAGGAATCGCTCAGCGTAAgtttaagagctatgc |
| BraALA3a-sgRNA07-F | CTACTGATCCTTGCGGAAAT |
| BraALA3a-sgRNA07-R | CTCGCAGAACAAAGACCTAG |
| TF-BraALA3a-sgRNA07 | ctcggagtgatcgcacTCTGATAAGACTGGAACGCTGAC |
| TR-BraALA3a-sgRNA07 | ctgagaggctggatggGAGAGACAAATTAGACATGCTTCC |
| Crispr-BraALA3a-sg03-F | cctctaatacgactcactataGGAAAGGCCCTTGAAAGGACAgtttaagagctatgc |
| BraALA3a-sgRNA03-F | AGGTGCTTCTTCTTTGACTTCC |
| BraALA3a-sgRNA03-R | ATGAGCCATAACAACTCTTTCC |
| pCA-Cas9-F | CAAGTACGTGAACTTCCTCTACC |
| pCA-Cas9-R | GCTGGGAAAGGTCGATACGAGTC |
| Bar-F | ATGAGCCCAGAACGACGCCCGGC |
| Bar-R | TTAAATCTCGGTGACGGGCAGGACC |
| **Genetic transformation** | |
| 1300-BraALA3a-GFP-F | agaacacgggggacgagctcATGGTTCGATCGGGTAATTTAAGCG |
| 1300-BraALA3a-GFP-R | ctagaggatccccgggtaccCTTCTTCTTTGGTACTGTCGGCC |
| 1300-BraALA3b-GFP-F | agaacacgggggacgagctcATGGTACGATCGGGTAGTTTAAACG |
| 1300-BraALA3b-GFP-R | ctagaggatccccgggtaccCTTCTTTGGTACTTTAGGGCGTGAC |
| 1300-BraALA3c-GFP-F | agaacacgggggacgagctcATGGCTAGCTCCGGAGGTGGATTAA |
| 1300-BraALA3c-GFP-R | ctagaggatccccgggtaccTTTCTTCTTTGGTGCTTTGGGTCGTG |
| HygB-F | TGTAGTGTATTGACCGATTCCTTGC |
| HygB-R | GTTCGACAGCGTCTCCGACCTGAT |

**Table S3.** SPR binding affinities between PCZ and BraALA3a/b/c homologous proteins.

| Compound | Protein | Avg ka (1/Ms) | Avg kd (1/s) | **Avg KD (M)** | Int.Intensity Level | ABS (tr_KD) |
|---|---|---|---|---|---|---|
| PCZ | BraALA3a | 2.05E+05 | 1.26E-02 | 6.14E-08 | Strong | 23.9564 |
| PCZ | BraALA3b | 1.71E+03 | 1.80E-02 | 1.05E-05 | Middle | 16.5389 |
| PCZ | BraALA3c | 4.57E+04 | 6.41E-03 | 1.40E-07 | Strong | 22.7649 |
| DMSO | BraALA3a | 1.26E+00 | 7.01E-01 | 5.58E-01 | VW/None | 0.8418 |
| DMSO | BraALA3b | 1.37E+00 | 7.29E-01 | 5.30E-01 | VW/None | 0.9146 |
| DMSO | BraALA3c | 2.15E+00 | 7.75E-01 | 3.60E-01 | VW/None | 1.4758 |

SPR affinity coefficients of BraALA3a/b/c-BraALIS2 and PCZ. Avg Ka (1/Ms): The average association rate constant (Ka) represents the ratio of complex formation per unit time relative to initial reactant concentrations, where higher values indicate faster molecular binding. Avg Kd (1/s): The mean dissociation rate constant (Kd) reflects the proportion of complex dissociation per unit time, with elevated values corresponding to faster complex disintegration. Avg KD (M): The equilibrium dissociation constant (KD = Kd/Ka) quantifies binding affinity at dynamic equilibrium. Lower KD values signify stronger intermolecular interactions. Interaction Intensity Level: Determination of affinity. KD range: $10^{-13}$-$10^{-5}$ M = strong; $10^{-5}$-$10^{-3}$ M = moderate; $10^{-3}$-$2\times10^{-2}$ M = weak; >$2\times10^{-2}$ M = negligible. ABS (tr_KD)：Absolute affinity coefficient. ABS (tr_KD) = ABS ($\log_2$KD), where higher numerical values indicate enhanced binding affinity. Reported as mean ± SD from four technical replicates.

**Method S1.** Measurement of PCZ level by UPLC-MS/MS.

PCZ-treated flowering Chinese cabbage samples (10 g) were flash-frozen in liquid nitrogen, homogenized by grinding, subjected to ultrasonication (20 mL grade acetonitrile，4 g MgSO4，1 g NaAc) and centrifugation (4,500 rpm, 10 min), then processed through solid-phase extraction (75 mg PSA，425 mg $MgSO_4$，50 mg C18 adsorbent) and filtration (0.22 μm) before UPLC-MS/MS analysis. Each experimental run included 3 technical replicates, with each replicate pool comprising 10 seedlings.

Chromatographic separation was performed on a Waters ACQUITY UPLC BEH C18 column (2.1 × 100 mm, 1.7 μm) maintained at 40°C, with mobile phase A (0.1% formic acid in water) and B (acetonitrile) under the following gradient: 10% B (0-2 min), 10%-95% B (2-5 min), 95% B (5-7 min), returning to 10% B at 7.1 min. The flow rate was 0.3 mL/min with 1 μL injection volume.

Mass spectrometric analysis was conducted using electrospray ionization (ESI) in positive-ion multiple reaction monitoring (MRM) mode, with the following parameters: nebulizing gas (3 L/min), heating gas (10 L/min), drying gas (10 L/min), interface temperature (300°C), desolvation line temperature (250°C), and block heater temperature (400°C). The capillary voltage was maintained at 2.5 kV with a cone voltage of 15 V. PCZ eluted at 3.42 min, monitored via qualifier ion transition 342.0/69.0 (collision energy: 22 eV) and quantifier transition 342.0/159.0 (collision energy: 30 eV).[7]

**Method S2.** Histological section.

PCZ-treated flowering Chinese cabbage samples were fixed overnight at 4°C in fixative (2.5% Glutaric dialdehyde, 0.1 M Phosphate Buffer (pH7.2), 0.1% Tween-20 and 4% Paraformaldehyde), followed by graded ethanol dehydration. The samples were treated with a mixture of acetone and embedding agent with V/V = 3/1, 2/1, 1/1, 1/2, 1/2 and 1/3 for 2 h. Finally, the samples were treated twice with pure embedding agent for 24 h each time. The samples were embedded by Eponate 12™–Araldite embedding Kit with DMP-30 (TED PELLA, INC). The samples were polymerized at 60°C for 16-24 h until the samples cooled to certain hardness. The samples were trimmed to the appropriate size and section, embedded plant tissues were sectioned by the *Leica RM2235* manual microtome in 5-10 μm. The toluidine blue-stained sections were observed under Laser capture micro-dissection (LMD).[27] Cell length and width were quantified from stained samples using Image-Pro Plus 6.0 software.

**Method S3.** Heterologous expression in yeast.

Total RNA of flowering Chinese cabbage was extracted with the E.Z.N.A. Plant RNA Kit (Omega, Guangzhou, China), and reverse transcription was conducted with the PrimeScript™ RT reagent Kit and gDNA Eraser (Takara, Dalian, China). The CDS region of *BraALA3a/b/c* was amplified by 2 × Primer STAR Max premix (Takara), *BraALA3a/b/c* was subcloned into pYES2 by ClonExpression®II One Step Cloning Kit (Vazyme, Nanjing, China). Primers were shown in Table S3. The pYES2-*BraALA3* were supplemented in *drs2* yeast or BY4741 (*MATa his3-1 leu2-0 met15-0 ura3-0*). The preparation and transformation of yeast receptive state were carried out according to Yeastmaker™ Yeast Transformation System 2 (Takara), and positive transformants were identified using Quick Yeast positive clone assay Kit (Coolaber, Beijing, China).

Complement transformants and enriched expression transformants were incubated in 500 μL SD/-His-Leu-Met medium (0.67% YNB, 2% D-Galactose, 0.002% His, 0.002% Met, 0.01% Leu, pH5.7) for 12 h at 210 rpm, 30°C. Then, transformants were replaced with fresh culture medium and cultured for 4 h, $OD_{600}$ was adjusted to 0.4, 0.04 (1: 10), 0.002 (1: 50) and 0.004 (1: 100) with distilled water ($ddH_2O$). 3 μL transformants diluted fluid were spotted onto SD/-His-Leu-Met medium (2% Agar) containing DMSO, 0.5 or 1.0 μM PCZ (or other compounds), and cultured at 30°C for 2-3 d.[30]

Complement transformants and enriched expression transformants were incubated in 3 mL SD/-His-Leu-Met medium overnight at 30°C. $OD_{600}$ was adjusted to 0.04 with corresponding fresh culture medium and DMSO/0.5 μM PCZ. Yeast cells were induced to express at 210 rpm, 30°C, and $OD_{600}$ was measured at 9, 12, 15, 18, 21, 24, 33, 36, 39, 42, 45, 48, 60 and 72 h for growth curve plotting.[27,29]

**Method S4.** Generation of *BraALA3a/b/c*$^{atala3}$ complemented lines.

*A. thaliana* transformation was performed via floral dip using GV3101 harboring the 1300-BraALA3a/b/c-GFP construct.[31] Transformant were cultured in LB medium (10 g/L Tryptone，5 g/L Yeast extract，10 g/L NaCl) with antibiotics (50 μg/mL Kan，50 μg/mL Str，20 μg/mL Rif), and resuspended in infiltration medium (5% Sucrose，0.02% SilwetL-77，$OD_{600}$ = 1.0), *A. thaliana atala3* mutant inflorescences were immersed for 30 s, followed by 24 h dark incubation before returning to standard growth conditions until silique maturation. Transgenic progeny was selected by Hyg resistance and confirmed through molecular genotyping.

**Method S5.** Molecular docking of BraALA3a/b/c-BraALIS2 with PCZ.

The conformational structures of DRS2-Cdc50p protein were obtained from Protein Data

Bank.[22] The amino acid sequences of the proteins were analysed for physicochemical properties including parameters such as molecular weight, theoretical isoelectric point, and total atomic number using ProtParam tool. The 3D structures of the ligand molecules PCZ, and PI4P were downloaded from the PubChem database. High-resolution crystal structures have not been reported for BraALA3a/b/c-BraALIS2 double-stranded proteins, and the artificial intelligence model ColabFold (https://colabfold.mmseqs.com/) was used for protein 3D structure modelling.[32] The amino acid structures were obtained from the UniProt database, and were hosted on the Google Colaboratory project hosting platform using ColabFold, the structure of the polypeptide chain complexes was predicted by multichain mode, with other parameters defaulted. After submitting the task, ColabFold was trained using transformer model and outputed a predicted structure file (PDB format). The predicted model was reviewed by PyMOL visualisation software. The complex model was evaluated and optimised using the methods of Ramachandran.[33]

Water molecules and other small-molecule ligands in the protein structure were removed using PyMOL to check the protein atom type and structural integrity, and then the protein molecules were added with polar hydrogen atoms and Gaussian charges and the ligand small molecules were set up with torsionally reversible bonding positions using AutoDock Tools to obtain a standard PDBQT file. Based on previous reports,[22,34] the parameters of the docking box were determined (box dimensions: 30 Å × 30 Å × 30 Å, lattice length: 0.375 Å). Molecular docking was performed using Autodock Vina,[35,36] vina.exe was used for docking and vina_split.exe was used to split the docking results, exhaustiveness = 100 was setted for more accurate results. The algorithm of Autodock Vina was set as lazy scoring function and genetic algorithm when executing docking to find the best binding conformation and affinity between ligand and receptor in the set search space. The ligand-receptor interactions in each binding mode, including hydrogen bonding, hydrophobic forces, and van der Waals forces between the ligand and active site residues, were analysed by PyMOL and UCSF ChimeraX.

**Method S6.** Molecular dynamics simulation (MD).

50 ns all-atom molecular dynamics simulations of the complex model were performed using the GROMACS software package.[37] The structure files of proteins and small molecule ligands were uploaded through the Protein/Ligand Complex module of the CHARMM-GUI online server (https://charmm-gui.org/), and the structures were processed for centre of mass alignment, residue error correction, and missing atom construction, and the system was run through the process of ionisation (0.15 mol/L NaCl) and solvation (TIP3P water model) to

construct topology files. Energy minimisation files were generated using the gmx grompp tool of GROMACS, with the maximum number of steps set to 50,000, and conjugate gradient energy minimisation was carried out via the gmx mdrun command with a time step of 1 fs. The NPT equilibrium condition was constant temperature (T = 300 K) and constant pressure (P = 1 bar), and was performed using a V-rescale temperature coupling and a Parrinello-Rahman pressure coupling, with a simulation duration of 100 picoseconds. Molecular dynamics simulations of final conformation production based on NPT equilibrium were performed with gmx grompp to generate input files for 50 ns molecular dynamics production runs. The final molecular dynamics trajectories were used to analyse the structure, conformational dynamics and interaction characteristics of protein-small molecule complexes. The stability of conformations was assessed by Root Mean Square Deviation (RMSD), Root Mean Square Fluctuation (RMSF), Radius of Gyration (Rg), Solvent-Accessible Surface Area (SASA) and number of hydrogen bonds.[38]

**Method S7.** P4-ATPase protein expression.

The engineered *Saccharomyces cerevisiae* strain BJ5465 (a high-yield protein expression system with multiple auxotrophic markers) was utilized for P4-ATPase. BraALA3a/b/c and *β*-subunits from flowering Chinese cabbage were cloned into pRS426 (sfGFP-TwinStrep-3C-CtP4-ATPase, Ura resistance selection) and pRS424 (sfGFP-His-3C-β-subunit, Trp resistance selection) vectors, respectively, then co-transformed into BJ5465 (*MATa ura3-52 trp1 leu2-delta1 his3-delta200 pep4::HIS3 prb1-delta1.6R can1 GAL*). Transformants were selected on SD-Ura/Trp medium (6.7 g/L YNB with ammonium sulfate, 1.29 g/L DO supplement-Ura/Trp, 2% glucose, pH5.7) and verified by colony PCR. Selected clones were cultured in SD-Ura/Trp with 2% raffinose (30°C, 220 rpm, 24 h) until $OD^{600}$ reached 5, then induced with 4×YPD medium (40 g/L yeast extract, 80 g/L peptone, 8% galactose) at 25°C, 220 rpm, 20 h. Cells were harvested by centrifugation (4°C, 3,000 × g, 20 min) for protein extraction.

Membrane proteins were extracted through differential centrifugation and detergent solubilization. Yeast cells were resuspended in 2 volumes of lysis buffer (20 mM Tris-HCl [pH7.4], 150 mM NaCl, 5 mM $MgCl_2$, 1 mM DTT, 1:500 protease inhibitor cocktail), disrupted by high-pressure homogenization (1,800 bar, 5-7 passes), and clarified by centrifugation (4°C, 20,000 × g, 25 min). Membrane fractions were collected via ultracentrifugation ((4°C, 44,000 rpm, 1 h, 2 passes), then solubilized in 5 volumes of extraction buffer (20 mM Tris-HCl (pH7.4), 10% (v/v) glycerol, 150 mM NaCl, 5 mM $MgCl_2$, 2% lauryl maltose neopentyl glycol (LMNG), 1 mM DTT, 1:500 protease inhibitor cocktail) for 1 h at 4°C. The solubilized fraction was recovered by ultracentrifugation (4°C, 44,000 rpm, 1 h) for subsequent analyses.[39] Protein expression and purity were assessed by SDS-PAGE, with concentration determined using Enhanced BCA Protein Assay Kit (Beyotime, Shanghai, China).

**Method S8.** Binding affinity determination by SPR.

Protein-ligand interactions between BraALA3a/b/c-BraALIS2 dimers and PCZ were analyzed by surface plasmon resonance (SPR). Compounds were immobilized on 3D photo-crosslinked chips using a Biodot™ AD1520 array spotter, followed by vacuum drying, UV crosslinking, and sequential washing (DMF, ethanol, $H_2O$; 15 min each) before nitrogen drying and Flowcell Cover assembly. Detergent/glycerol-free protein samples (PBS-dialyzed and concentrated) were diluted, and diluted in PBST (pH7.4, 0.1% Tween 20) at five concentrations (10, 40, 160, 640, 2560 nM). Samples were injected in ascending concentration order (0.5 µL/s, 4°C, 600 s association/360 s dissociation), with regeneration using glycine-HCl (pH 2.0, 2 µL/s). Kinetic parameters (Ka, Kd) and equilibrium dissociation constants (KD) were derived from binding curves to quantify interaction strengths.[40]

**Method S9.** *In vitro* ATPase activity assay.

Functional validation of protein complex bioactivity was performed through in vitro ATPase activity assays, including testing for potential P4-ATPase activators or inhibitors.[39] Using a Malachite Green Phosphate Assay Kit (Sigma), liberated phosphate from ATP hydrolysis was quantified colorimetrically at 620 nm with or without phosphatidylserine (POPS), with standard calibration curves enabling precise activity quantification across experimental conditions. POPS was prepared as 10 mM stock in 1% $C_{12}E_9$ solution, while ATP (12.5 mM stock) was aliquoted and stored at -80°C. Protein complexes were pre-incubated with PCZ for 1 h prior to ATP addition, with $BeF3^-$ (a known P-type ATPase inhibitor) serving as negative control. All experimental conditions were independently replicated three times.

**Method S10.** Quantitative real-time PCR (qRT-PCR).

Total RNA was extracted from PCZ-treated flowering Chinese cabbage organs followed by reverse transcription. RT-qPCR was performed using ChamQ SYBR qPCR Master Mix (Vazyme, Nanjing, China). *UBC10* served as the reference gene for flowering Chinese cabbage. All experiments were performed with 3 independent biological replicates. The Ct value from the BioRad CFX96TM Real Time PCR System were derived,[41,42] the expression level of the target genes was calculated under different treatments using equation $2^{-\Delta\Delta Ct}$ (the expression level of the internal reference gene is 1). The primers used for RT-qPCR analysis are listed in

Table S3.

**Method S11.** Generation of *BraALA3a/b/c*-overexpressing lines in flowering Chinese cabbage. *Agrobacterium*-mediated transformation of flowering Chinese cabbage was performed by cloning *BraALA3a/b/c* coding sequences into the pBI121 binary vector, followed by GV3101 transformation and cultured in LB medium (10 g/L tryptone, 5 g/L yeast extract, 10 g/L NaCl) with antibiotics (50 μg/mL Kan，20 μg/mL Rif). Cotyledon-petiole explants from in vitro-grown seedlings (1/2 MS，3%Sucrose，0.75%Agar，pH5.8) underwent pre-culture (MS，3%Sucrose，0.75%Agar，pH 5.8，2 mg/L trans-Zeatin (TZ)，0.5 mg/L IAA，7 mg/L $AgNO_3$), bacterial infection (MS，3%Sucrose，pH 5.8，100 μM Acetosyringone (AS)), co-cultivation (MS，3%Sucrose，0.75%Agar，pH 5.8，2 mg/L TZ，0.5 mg/L IAA，7 mg/L $AgNO_3$，100 μM AS), antibacterial differentiation (MS，3%Sucrose，0.75%Agar，pH 5.8，2 mg/L TZ，0.5 mg/L IAA，7 mg/L $AgNO_3$，300 mg/L Timentin (TMT)), antibiotic selection (MS，3%Sucrose，0.75%Agar，pH 5.8，2 mg/L TZ，0.5 mg/L IAA，7 mg/L $AgNO_3$，300 mg/L TMT，10 mg/L Neomycin [Neo]), shoot regeneration (MS，3%Sucrose，0.75%Agar，pH 5.8，2 mg/L TZ ，0.5 mg/L IAA，7 mg/L $AgNO_3$，300 mg/L TMT), root regeneration (MS，3%Sucrose，0.75%Agar，pH 5.8，0.3 mg/L IAA，300 mg/L TMT),[5] and acclimatization to generate *BraALA3a/b/c*-overexpressing plants. Transgenic lines were screened by PCR and selected for high transgene expression via qRT-PCR.

**Method S12.** Generation of *BraALA3a/b/c* knockdown lines in flowering Chinese cabbage. Artificial microRNA (amiRNA) sequences targeting *BraALA3a/b/c* were designed using WMD3 (http://wmd3.weigelworld.org), validated for specificity via psRNATarget (http://www.plantgrn.org/psRNATarget/), and assessed for secondary structure using UNAFold (http://www.unafold.rna.albany.edu). Synthesized fragments (Beijing Tsingke Biotech) were cloned downstream of the CaMV 35S promoter in pBI121 via homologous recombination. The construct was transformed into Agrobacterium tumefaciens GV3101, cultured in LB medium (10 g/L tryptone, 5 g/L yeast extract, 10 g/L NaCl) with antibiotics (50 μg/mL Kan, 50 μg/mL streptomycin [Str], 20 μg/mL rifampicin [Rif]). Cells were harvested by centrifugation and resuspended in infiltration buffer (5% sucrose, 0.02% Silwet L-77, $OD_{600}$ = 1.5). flowering Chinese cabbage 'Youqing Sijiu' was transformed via vacuum-infiltration floral dip (-0.09 MPa), with 6-8 manual pollinations using non-transformed pollen. After removing secondary bolts, T1 seeds were harvested at 8-10 weeks. Transgenic plants were screened by PCR and validated by qRT-PCR.[42]

**Method S13.** Physiological parameter quantification in flowering Chinese cabbage.

Soluble protein content was determined by coomassie brilliant blue assay. Fresh aerial tissues (0.5 g) were ground in liquid nitrogen, homogenized in distilled water, mixed with 5 mL Coomassie Brilliant Blue G-250 reagent, and incubated for 2 min. and measured at 595 nm absorbance, with concentrations calculated against standard curve.

Reduced vitamin C content was determined by molybdenum blue assay. Frozen tissue (1 g) was homogenized in liquid nitrogen, extracted with 10 mL oxalic acid-EDTA solution, diluted to 50 mL, and filtered. 10 mL aliquot was mixed with 1 mL metaphosphoric-acetic acid solution, 2 mL 5% sulfuric acid, and 4 mL ammonium molybdate reagent, incubated for 15 min, and measured at 709 nm absorbance.

Soluble sugar content was determined by anthrone assay. Frozen tissue (0.5 g) was homogenized in liquid nitrogen, extracted with 10 mL distilled water at 100°C for 1 h, then filtered after cooling. 0.1 mL aliquot was mixed with 1.9 mL water, 0.5 mL ethyl anthrone reagent, and 5 mL concentrated sulfuric acid, vortexed vigorously, and measured at 630 nm absorbance, with concentrations calculated against standard curve.

Chlorophyll content was determined by acetone/ethanol assay. Fresh leaf tissue (0.5 g) was immersed in 25 mL of acetone/ethanol (1:1, v/v), incubated in darkness for 16 h, and the supernatant was analyzed for chlorophyll A (663 nm) and chlorophyll B (645 nm) absorbance. All measurements included three biological replicates.[2,5]

**Method S14.** Establishment of a rapid screening system based on protoplast for gene-editing targets in flowering Chinese cabbage.

High-efficiency sgRNAs targeting *BraALA3* were designed using CRISPR-GE (http://skl.scau.edu.cn/) with evaluation of off-target potential, followed by in vitro synthesis and transcription using the Guide-it sgRNA In Vitro Transcription System (Takara). Cleavage efficiency was validated through in vitro DNA digestion assays.

Five-day-old flowering Chinese cabbage seedlings were enzymatically digested in protoplast isolation solution (1.5% Cellulase R10，0.25% Macerozyme R10，20 mM MES (pH 5.7), 0.8 M mannitol，20 mM KCl，10 mM $CaCl_2 \cdot 2H_2O$，0.1% BSA，0.22 μm filter for sterilization), 25°C, darkness, 50 rpm, 3.5 h. The digestate was filtered through 40 μm meshes, centrifuged (60 × g, 3 min), washed with W5 solution (154 mM NaCl，125 mM $CaCl_2 \cdot 2H_2O$，5 mM KCl，2 mM MES (pH 5.7), 5 mM Glucose), and assessed for cell viability using fluorescein diacetate (FDA) staining under microscopy. Prior to transfection, cells were resuspended in MMG solution (0.4 M mannitol，15 mM $MgCl_2$，4 mM MES (pH 5.7)) at optimal density ($2 \times 10^5$ cells/g FW).

RNP complexes were assembled by mixing Cas9 protein and sgRNA at a 5:1 mass ratio, then delivered into flowering Chinese cabbage protoplasts via PEG-mediated transfection (40% PEG 4000， 0.2 M mannitol， 0.1 M $CaCl_2 \cdot 2H_2O$) with 10 min incubation at 25°C. After W5 solution washing, transfected protoplasts were cultured in 1N0.3K medium (1/2 MS， 3% sucrose， 0.4 M mannitol， pH 5.7， 1 mg/L NAA， 0.3 mg/L kinetin) for 14 days.[43,44] Genomic DNA was extracted using the MicroElute Genomic DNA Kit (Omega), with target sites amplified for RNP cleavage validation. Mutation profiles were characterized by Sanger sequencing or high-throughput sequencing,[45] confirming editable targets.
